# Lethal permeabilization of host bacteria to small-molecule compounds during phage penetration

**DOI:** 10.64898/2026.01.24.701513

**Authors:** Zihao Yu, Chaohua Wu, Harrison R. Lee, Ian Mattingly, Lily Torrans, Ryland F. Young, Lanying Zeng

## Abstract

The tailed phages (Caudoviricetes) rely on specialized mechanisms to inject their double-stranded linear genomic DNA (gDNA) across the bacterial envelope. During this gDNA entry process, ionic fluxes and membrane depolarization may occur, potentially leading to increased vulnerability to antibiotic molecules. Here, we report that phage gDNA translocation combined with certain small molecules triggers a rapid and phage-specific cell permeabilization leading to growth arrest. Using live-cell fluorescence microscopy, we visualized the accumulation of otherwise excluded compounds such as anthracyclines and propidium iodide. We show that the influx and accumulation of these compounds depend on both the presence of an inner membrane (IM) receptor and persistent membrane depolarization. We further demonstrate that phage-induced permeabilization enhances the efficacy of certain antibiotics, enabling a synergistic elimination of antibiotic-resistant bacteria. Our results provide a mechanistic foundation for combining phages with small-molecule therapeutics and offer new insights into how phage entry relies on and alters host physiology.

---

When infecting Gram-negative bacteria, phage genomic material (gDNA or gRNA) must penetrate both outer (OM) and inner membranes (IM) of the cell ^1^. To facilitate the penetration (for simplicity, the terms penetration and injection are used interchangeably unless otherwise specified), phages use diverse host factors such as lipopolysaccharides (LPS), porins, surface appendages, sugar transporters, periplasmic chaperones, and RNA polymerase ^2,3^. The penetration process is often accompanied by increased permeability, evident from the reversible and irreversible loss of cellular content, of which the extent and duration vary widely among phages ^4–8^.

In the case of tailed phages (Caudoviricetes), the injection mechanism and associated ionic fluxes generally correlate with the tail morphotypes. For example, siphophages like T1 and λ have a non-contractile tail and rely on separate receptors in the OM and IM for gDNA penetration, during which a substantial potassium ion efflux has been observed ^9,10^. Another siphophage, coliphage T5, follows a unique two-step transfer mechanism, where potassium efflux occurs during DNA translocation steps, but not during the pause between these two steps in the injection process ^11^. In contrast, myophages like T4 utilize tail contraction to facilitate gDNA translocation. During this process, transient ion fluxes stop once DNA internalization is complete. Indeed, infection with ghost phages (phages without gDNA) results in unabated ion release ^12,13^. In contrast, some podophages, such as the coliphages T3 and T7, accomplish gDNA penetration without detectable ion leakage ^14^. In these infections, a complex spanning the entire envelope is formed and the gDNA is pulled into the cytoplasm by RNA polymerase (RNAP) ^15^. In addition, phages often encode peptidoglycan hydrolase activities within the tail structure or as ejected proteins ^16^, adding an additional layer of cell envelope stress and disruption.

The molecular machines that drive phage genome delivery across two membranes remain poorly understood ^1,14^. Ionic fluxes associated with genome delivery appear to occur in both directions: phage infection induces the influx of H^+^ and Na^+^ into the cells, while potassium efflux results in either the partial or complete loss of total intracellular K^+^, as observed in T-phages and lambda infections ^5,9,10,17^. The nature and dynamics of the channel, as well as the specific membrane damage that underlies these flows, remain unclear. Moreover, it is not known how the gDNA penetration process comes to an end to close the channel and re-energize the membrane.

To address these questions and gain a deeper understanding of the critical step of phage infection, we seek to develop a method to quantitatively track gDNA penetration and monitor permeability changes using live-cell microscopy. We considered using naturally occurring secondary metabolites with intrinsic fluorescence to visualize and track the passage of substances across the envelope. *Streptomyces* species, well-known for their metabolic diversity, produce a wide range of secondary metabolites, including more than two-thirds of clinically relevant antibiotics ^18^. A previous study found that secondary metabolites produced by *Streptomyces* species exhibit inhibitory effects on phage propagation ^19^. Purified small-molecule compounds from the spent medium of *Streptomyces* spp. cultures were screened with various bacterium-phage systems, showing strong inhibition function against a range of phages infecting *E. coli* and *P. aeruginosa*. One notable group of these secondary metabolites that showed a strong inhibitory effect is the anthracyclines, including daunorubicin, doxorubicin, epirubicin, and idarubicin, many of which are commonly used in cancer chemotherapy ^20,21^. In fact, the inhibitory function of daunorubicin (daunomycin) on coliphage propagation was first reported more than half a century ago ^22^. Both studies demonstrated that the degree of inhibition varied across a panel of different phages, and for certain types, phage production was unaffected. It was suggested that many of these compounds, which are known to function as DNA or RNA intercalators, preferentially inhibited early steps of phage DNA transcription and replication while leaving host chromosomal replication undisturbed ^19^. More specifically, the previous study suggested that the effect could be related to the step of viral DNA circularization, and thus the nature of the incoming phage genome (i.e., dsDNA with either ssDNA cohesive ends or terminal redundancy) ^23^ may be determinative in the extent of infection interference.

In this study, by employing time-lapse microscopy to monitor the uptake of various fluorescent DNA intercalating compounds, we revealed a phage-mediated cell permeabilization, which allows small-molecule DNA intercalators to cross otherwise intact cellular barriers, leading to growth arrest. The results are discussed in terms of a new perspective on the potential synergy between phage and antibiotic therapeutics.

## Results

### Phage infection results in the apparent permeation of small-molecule compounds

To follow the process of phage DNA injection and examine the time window during which the phage becomes sensitive to the DNA intercalator, we employed time-lapse fluorescence microscopy to study phage penetration and replication in the presence of daunorubicin in real-time. Daunorubicin possesses an intrinsic fluorescence ^24^, enabling us to follow its fluxes and DNA intercalating activity. Using a previously developed reporter phage (λLZ2002) with the fluorescent repressor/operator system (FROS) to label phage lambda gDNA as well as the phage capsid ^25^, we were able to follow individual phage infection and their gDNA injection at the single-cell level.

Consistent with the previous study ^19^, we observed the entry of λ DNA, which did not replicate inside the cell, in the presence of daunorubicin (**Fig. 1a**). However, in contrast to the notion that the intercalation of the drug interfered with gDNA circularization and thus blocked the downstream transcriptional program ^19,26^, we observed that the infection caused a dramatic accumulation of daunorubicin that resulted in growth arrest and consequent abortive infection (**Fig. 1a, movie S1**). Without phage infection, daunorubicin does not affect cell growth or accumulate in the cell under normal growth conditions (**Extended Data Fig. 1**). By using a replication-defective phage (λLZ770), we confirmed that the daunorubicin permeabilization does not rely on phage replication (**Fig. 1b, c**). We also found the same cell permeabilization and high intracellular signals for a synthetic DNA intercalating agent, propidium iodide, upon lambda infection (**Fig. 1b, c, Supplementary Fig. 1**).

**Figure 1.**
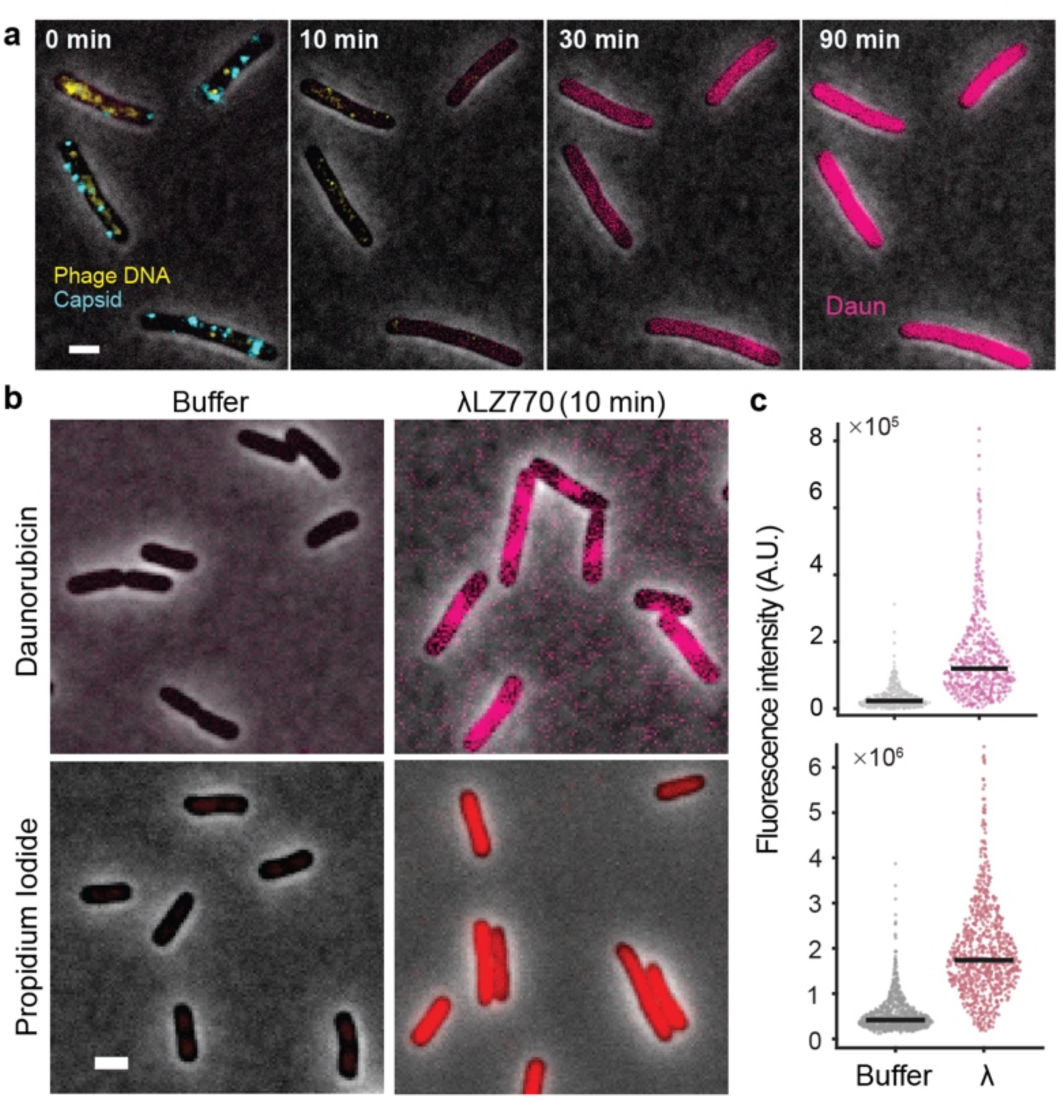
| Phage lambda infection leads to a compromised cell envelope and causes growth arrest with the addition of a DNA intercalating compound. **a)** Cells infected by lambda (cyan foci), as suggested by successful DNA injections (yellow foci), showed increasing daunorubicin uptake (magenta) and growth arrest. Cells were infected at an MOI of ∼5. Median filter applied to DNA and capsid channels for better visualization. Infection was performed in EZ-rich defined media. **b)** Cells infected by replication-defective mutant λLZ770 at an MOI of ∼5, in combination with 15 μM daunorubicin or 40 μM propidium iodide, show accumulation of these compounds after 10 min incubation at 30 °C. **c)** Quantification of intracellular signals in individual cells shown in **b**). Quantification results were pooled from three biological replicates. Black bars indicate the median of the dataset. Number of cells in the quantification: daunorubicin: buffer, 380 and phage, 653; propidium iodide: buffer, 902 and phage, 836. Scale bars = 2 µm.

### Cell permeabilization requires IM receptors

To understand the key steps during which the cell permeabilization may occur, we used IM receptor mutants to distinguish the model proposing that the cell leakage happens at either the OM, the IM, or both. Phage lambda requires an OM porin (maltoporin, LamB) as its primary receptor to trigger tail conformational changes and the initiation of genome ejection ^27,28^. On the other hand, lambda requires an IM receptor, the ManYZ complex, to achieve efficient gDNA translocation ^29,30^. While the exact mechanism of phage lambda gDNA translocation into the cytoplasm remains a mystery, it has been shown that most of the adsorbed phage particles on these *manY*/*manZ* mutants have the majority of their DNA remaining in the phage head, indicative of the defect in forming DNA translocating channels across the IM ^30^. When infecting *ΔmanY* and *ΔmanZ* strains (LZ3526 and LZ3527, respectively) with phage λLZ2002 in the presence of daunorubicin, we found that many fewer cells showed increased daunorubicin signals (**Fig. 2a, b)**, regardless of similar phage adsorptions (**Extended Data Fig. 2a-c**, and ^30^). Increased daunorubicin accumulation coincided with the successful entry of gDNA into the cytoplasm (**Fig. 2c, d**), and the overall fraction agrees well with the plating efficiency (**Extended Data Fig. 2d**). This indicates that cell permeabilization to daunorubicin depends on the successful penetration of the IM. We hypothesized that phage-induced damage to the IM could either lead to a large influx of the compound, resulting in cell death, or cause the normally transient process of membrane depolarization to become prolonged. The persistent disruption of the membrane potential thus promotes the entry of cell-impermeant compounds into the cell.

**Figure 2.**
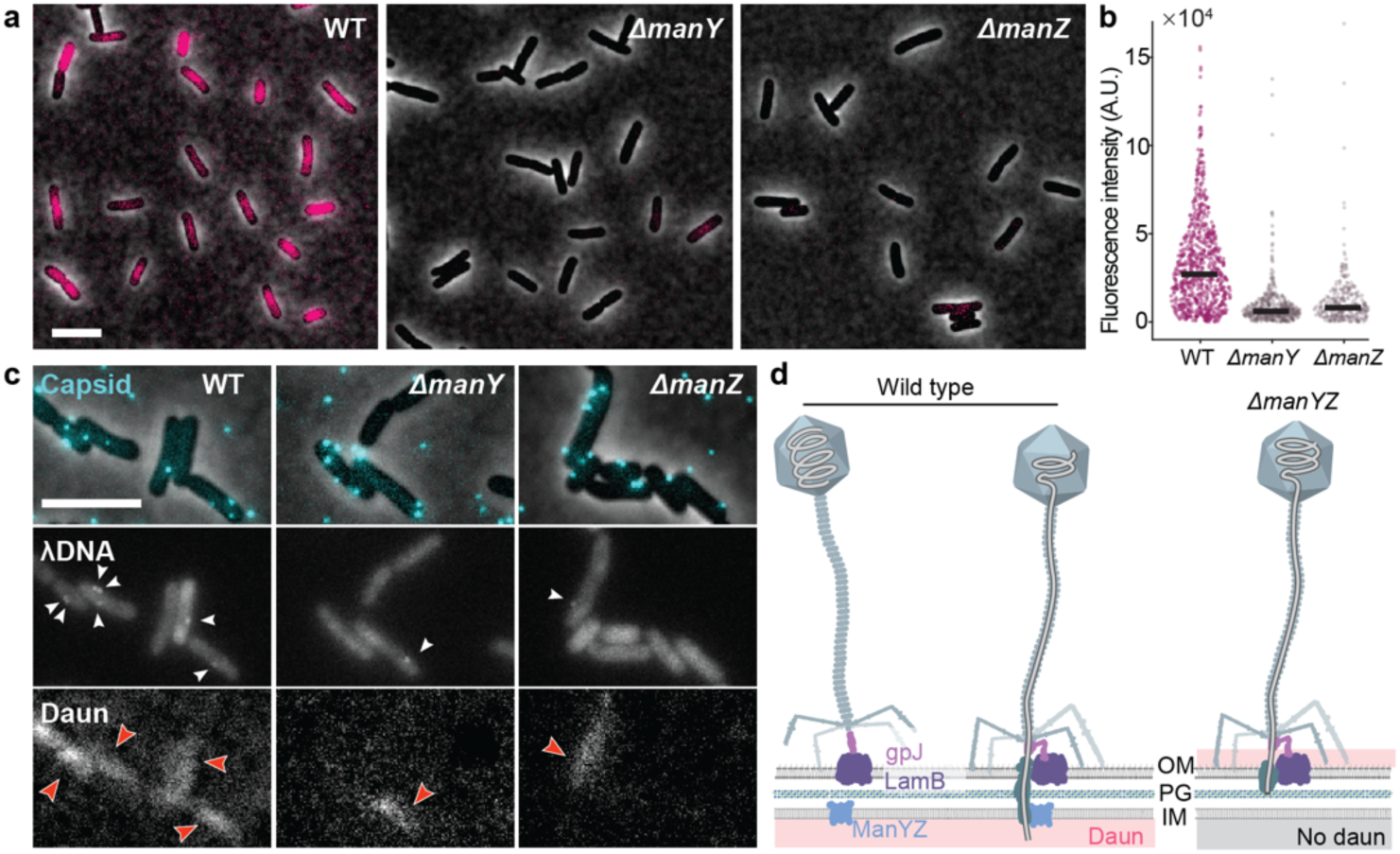
| The cell permeabilization by phage lambda is dependent on the IM receptor. **a)** Representative images of phage lambda-induced cell leakage to daunorubicin when infecting wildtype (LZ2001) and *ΔmanY*/*Z* mutants (LZ3526 and LZ3527) with λLZ2002. Deletion of either the *manY* or *manZ* gene results in a great reduction of intracellular signals. **b)** Quantitative analysis showing *ΔmanY/Z* mutants with less daunorubicin accumulation as exemplified in **a**). Number of cells used in this quantification: wildtype, 759; *ΔmanY*, 440; *ΔmanZ*, 325. **c)** Single-cell observation of phage adsorption (cyan foci) and DNA injection. DNA injection, indicated by the appearance of white foci (indicated by white arrowheads), is required for the accumulation of daunorubicin and the subsequent increase in intracellular fluorescence signals (red arrowheads). Since DNA injection is significantly reduced in the mutant strains (LZ3526 and LZ3527), these strains exhibit much weaker signals. To enhance visibility and facilitate comparison, the daunorubicin fluorescence contrast for the mutants was adjusted accordingly. **d)** Schematic illustration of lambda infecting WT compared to *ΔmanYZ* cells in terms of the formation of the IM channel for DNA ejection. The proposed model indicates that the trans-envelope channel is required for lambda-induced cell leakage to those small-molecule compounds. The adsorption of phage lambda at the outer membrane relies on the interaction between gpJ and LamB proteins, which is independent of the IM receptor ManYZ ^28,30^. Scale bars = 5 µm.

Irreversible adsorption of phage to the bacterial outer membrane triggers phage tail conformational change and creates a channel through OM, possibly compromising the OM barrier ^27,28^. To examine the function of OM as a barrier in limiting daunorubicin influx, we tested conditions that lead to an impaired outer membrane. We began by treating the cell with ethylenediaminetetraacetic acid (EDTA) to remove cations bound to LPS. This process weakens the lateral interactions between LPS molecules, thereby compromising the integrity of the outer membrane barrier ^31,32^. As expected, the cells appeared defective in growth and were often observed as filamented (**Extended Data Fig. 3a, b**); however, the intracellular fluorescence intensity showed only a small increase and did not exhibit a uniform pattern across the cells (**Extended Data Fig. 3b, c**), suggesting that a massive accumulation of daunorubicin did not occur, nor was the bacterial chromosomal DNA extensively bound by daunorubicin. A similar result was observed in the outer membrane hyperpermeable mutant *imp4213*, which exhibits sensitivity to organic compounds ^33^. This mutant permits the traverse of larger molecular-weight fluorescent dyes such as Atto488-ADA (M.W. = 674 g/mol, larger than 527 g/mol of daunorubicin), across the OM, enabling staining of the peptidoglycan (PG) layer ^34^. Microscopy revealed that, similarly to the EDTA treatment, the cells were filamented but did not exhibit strong intracellular daunorubicin signals (**Extended Data Fig. 4**). Based on this evidence, we concluded that the increasing OM permeability alone does not account for the phage-induced daunorubicin accumulation.

### Incomplete gDNA penetration of the IM is required for drug accumulation

Having found that the IM receptor ManYZ plays a key role during this process, we asked if the accumulation is directly related to the membrane potential. Membrane depolarization following phage infection has long been recognized ^4,35^. To confirm that the membrane depolarization can lead to daunorubicin accumulation, we treated the cells with the membrane potential-dissipating agent carbonyl cyanide m-chlorophenyl hydrazone (CCCP) in combination with daunorubicin. An accumulation of intracellular daunorubicin signals was observed (**Fig. 3a, Supplementary Fig. 2**). However, the phage-induced membrane depolarization should only be transient during a normal infection cycle, as re-energization of the IM is necessary for ATP production and hence successful phage propagation. We found that the cells were able to recover from membrane depolarization and remove intracellular daunorubicin after washes and resuspension in a CCCP-free fresh medium (**Fig. 3a, Supplementary Fig. 2**), even after a relatively long period of drug accumulation (30 min). Live-cell imaging of daunorubicin-accumulated cells in fresh LB showed a rapid removal of intracellular daunorubicin and recovered growth (**Fig. 3b**).

**Figure 3.**
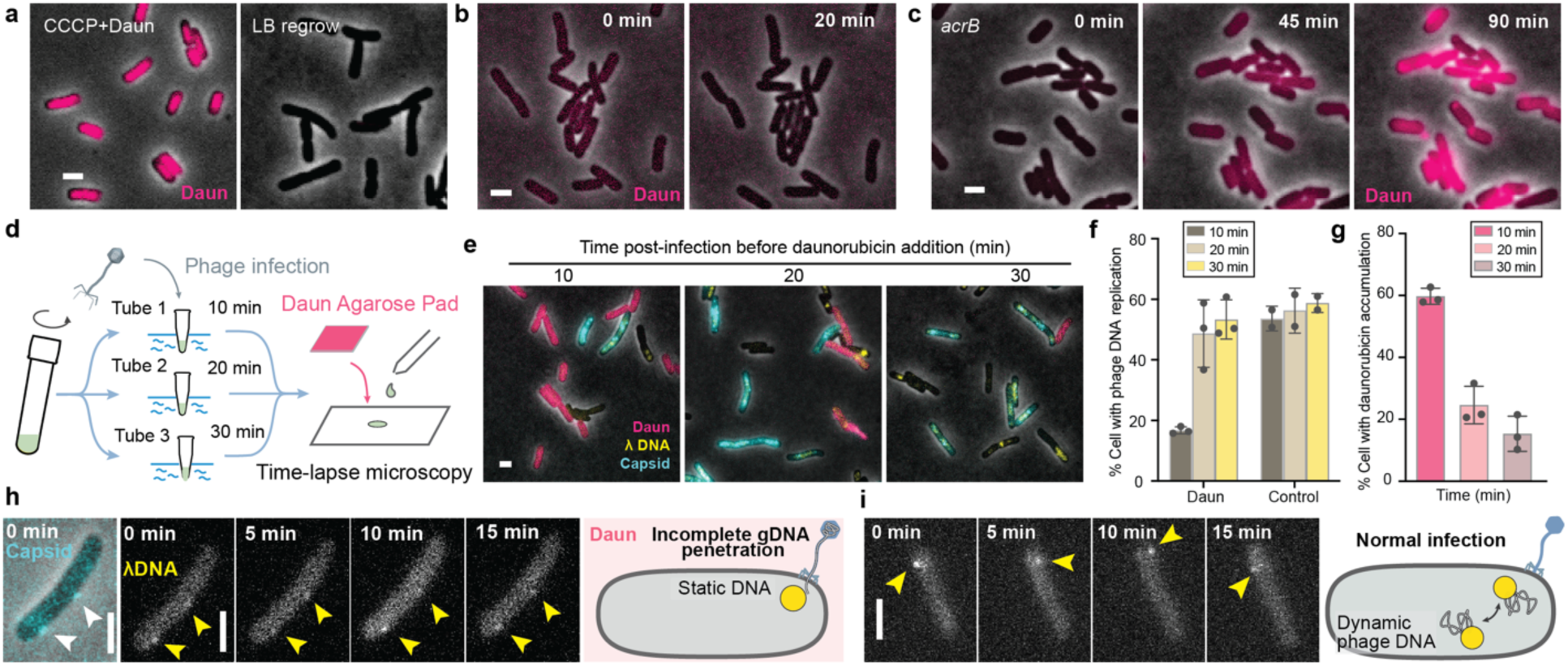
| Persistent disruption of the IM is required for drug accumulation. **a)** Representative images of daunorubicin accumulation in cells after CCCP treatment at room temperature for 15 min (left) and MG1655 cells can regrow in LB after removal of CCCP (right). **b)** Snapshots of MG1655 cells in the time-lapse movie to show a rapid removal of intracellular daunorubicin after CCCP treatment with daunorubicin and then regrowing in the LB agarose pad. **c)** *acrB* mutant cells show daunorubicin accumulation over time. **d)** Schematic of the procedure used to assay the transiency of permeabilization. **e)** Representative images at a late time point prior to lysis showing the accumulation of daunorubicin (magenta) or phage replication and assembly (capsid expression, cyan). When daunorubicin was introduced at a later time post-phage infection, less accumulation was observed in cells. **f)** Percentage of cells showing DNA replication, indicative of a normal phage life cycle without inhibition by daunorubicin. Control experiments were performed on LB agarose pads without daunorubicin. **g)** Percentage of cells showing daunorubicin accumulation and growth arrest. **h)** Phage DNA (yellow arrowheads) remains static near the cell surface and co-localizes with the capsid (white arrowheads) in the presence of daunorubicin during infection. This suggests incomplete gDNA penetration. **i)** Phage DNA moves freely in the cytosol in normal infection. Scale bars = 2 μm.

We reasoned that the reversible nature of the daunorubicin flux may be a result of the presence of a multidrug efflux pump, as suggested by previous studies of the existence of such an activity for another anthracycline, doxorubicin ^36^. When we disrupted the function of the AcrAB-TolC efflux system by knocking out the *acrB* gene, which encodes the IM component of the system, we found that the cells became sensitive to daunorubicin, exhibiting growth inhibition and an accumulation signal of the compound, similar to that induced by phage penetration (**Fig. 3c**). Thus, if the phage only causes transient membrane depolarization as happens during a normal infection cycle, the damage done by the transient depolarization would be soon overcome by the efflux system.

The transient nature of phage-induced disruption of the IM during a normal infection cycle is further supported by analyzing the time window during which cells are sensitive to drug treatment. Daunorubicin was added at various time points post-infection, and the ratio between productive lysis versus drug-dependent growth arrest was assessed using capsid-mTurquoise2 signals for phage production and red fluorescence for drug accumulation. Our data showed that cell sensitivity to daunorubicin decreased significantly from 10 minutes post-infection to 20 or 30 minutes (**Fig. 3d-g**). This suggests that the changes in permeability and membrane depolarization induced by the phage are transient, reverting to a less vulnerable state shortly after the completion of DNA penetration.

We hypothesized that the normal transient process becomes persistent through the following two layers of action. First, just as during normal infection, phage DNA injection causes an initial, transient disruption of the membrane potential and proton gradient which immediately impairs the activity of drug efflux pumps. Second, daunorubicin disrupts the gDNA translocation process and prevents the channel from resealing. This leaves a non-selective pore open, causing a constant leak of ions. Indeed, we found that the injected DNA in the presence of daunorubicin showed very restrained movement in the cytoplasm and was co-localized with the capsid for the full period of observation, which indicates incomplete gDNA injection (**Fig. 3h** and **Extended Data Fig. 5a**). In comparison, during regular infection without daunorubicin, we found that injected DNA showed dynamic motion in the cytoplasm (**Fig. 3i** and **Extended Data Fig. 5b, movie S2**).

### Infection-induced cell permeabilization is phage-specific

We have shown that lambda infection induces permeabilization to daunorubicin, leading to growth arrest. This provides an explanation for the reduced lambda fecundity in the presence of daunorubicin reported previously ^19,22^. However, what has also been shown in these studies is the variability in this effect among different phages. For example, coliphages T6 and T7 exhibit dramatically lower sensitivity to daunorubicin (six to seven orders of magnitude less than lambda).

With this in mind, we repeated our experiments with diverse phages in combination with daunorubicin. We first analyzed the accumulation of daunorubicin during infection by the canonical coliphages T4, T5, and T7. We found that phage-induced cell permeabilization by these phages differs drastically from that caused by phage lambda infections. Using time-lapse microscopy, we found that the T4-daunorubicin combination did not lead to an altered route of phage infection cycle: T4-infected cells underwent normal lysis and exhibited no detectable accumulation of daunorubicin before lysis (**Fig. 4a, movie S3**). Similar to T4, most T7 infections in the presence of daunorubicin also lead to successful lysis. However, a fraction of T7-infected cells (∼20%), although not taking up daunorubicin, undergo growth arrest and thus do not lyse (**Fig. 4a, movie S3**). The phenotype was also quantitatively validated by comparing the intracellular fluorescence signal at 10 min post-infection, where phage lysis has not yet occurred, and the effects of phage replication and phage gene expression are minimal. Compared to the daunorubicin accumulation caused by lambda infection, T4 and T7 infections led to a limited increase in intracellular daunorubicin signals at 10 min post-infection (**Fig. 4b**).

**Figure 4.**
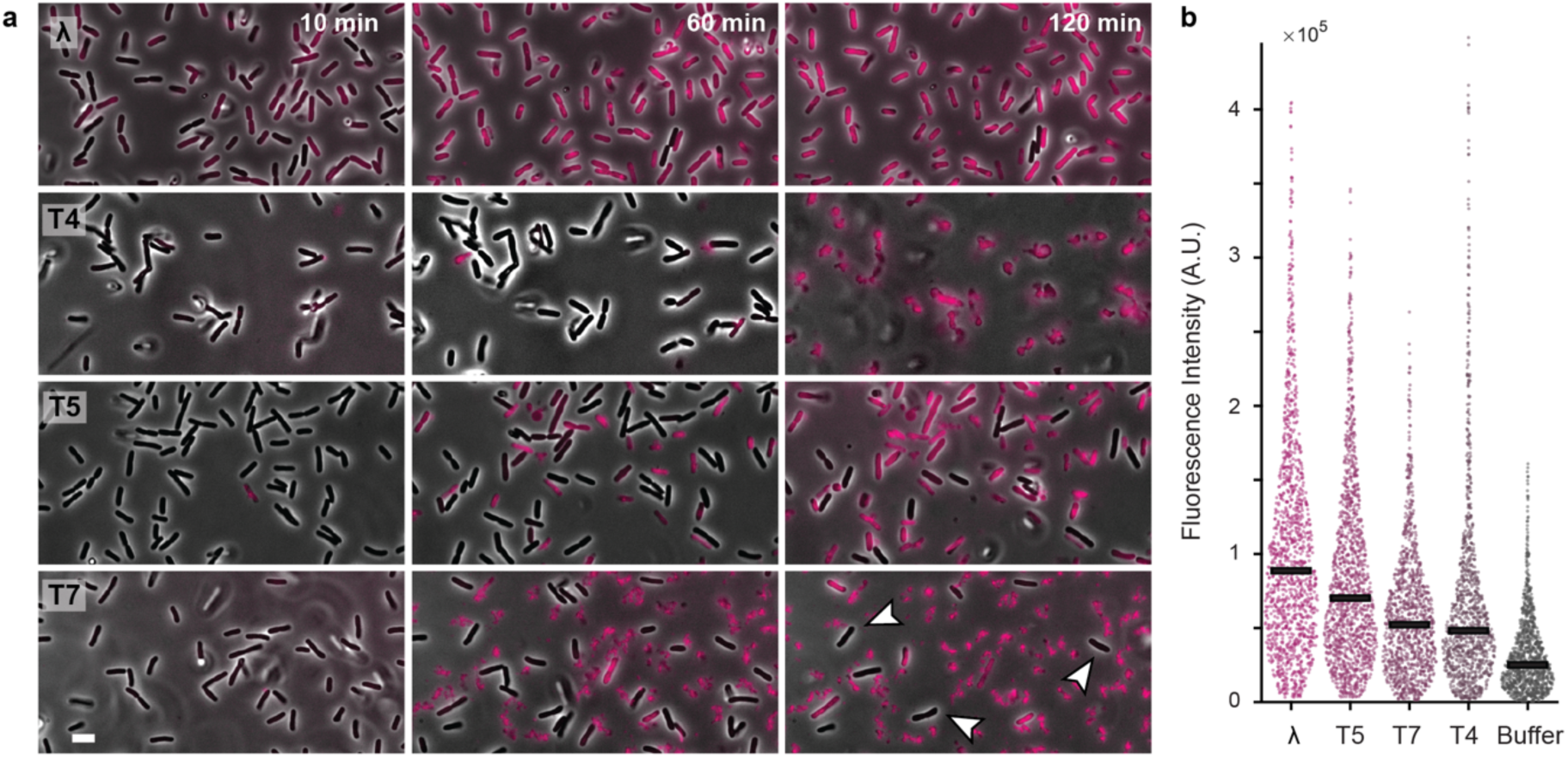
| Cell permeabilization to small-molecule compounds is phage-dependent. **a)** Representative snapshots from time-lapse movies of PL15 cells infected by λLZ613 and wildtype T-phages at high MOIs (>2), in the presence of daunorubicin. The cell permeability to daunorubicin differs dramatically among different phages. Note that at later time points, the daunorubicin fluorescence observed in T4- and T7-infected samples corresponds to cell lysis and dye bound to cell debris. **b)** Quantification of intracellular intensity early after treatments of different phages (10 min post-infection). Black bars indicate the median of the fluorescence intensity. Number of cells in the quantification: λ, 902; T5, 937; T7, 810; T4, 869; buffer, 683. Scale bar = 5 μm.

Phage T5, in combination with daunorubicin, leads to an abortive infection and growth arrest, similar to that caused by phage λ (**Fig. 4a**). However, it differs markedly from λ infection in that the infected cells appear physically compromised, likely because even partial genome internalization (e.g., the first-step transfer segment ^37^, or tail-associated fusogenic/lysozyme activities ^38^) exerts a more destructive effect on cell physiology, driving the cells beyond mere metabolic inhibition to become phase-light ghost cells (**Extended Data Fig. 6a, b**). The quantification of intracellular fluorescence signal at 10 min post-infection for T5-infected cells showed higher daunorubicin signals than T4 and T7 but is lower than lambda (**Fig. 4b**). This result was consistent with the inhibitory effect of daunorubicin at this concentration (15 μM) on the propagation of these phages in liquid cultures (**Extended Data Fig. 6f**), which is also consistent with the previous study ^19^ using a higher concentration of daunorubicin, further validating that the phage-induced permeabilization change is the cause of the phage production level changes.

A similar result was found when combining these T-phages with the synthetic DNA intercalator propidium iodide, both at the single-cell fluorescence level and at the bulk level by measuring productive phage production in liquid culture with propidium iodide (**Extended Data Fig. 6c-e, movie S3**). Overall, these data suggest a model in which the compound interferes with the phage penetration process across the two membranes, often leading to cell permeabilization, altered cellular physiology, and growth arrest, which correlate with reduced phage propagation in liquid culture.

### Compound-induced disruption of the phage injection channel enhances antibiotic activity

Above, we showed that certain phage infections, when combined with DNA intercalators, result in a disrupted injection process. This frequently leads to incomplete phage gDNA translocation and persistent IM leakage, ultimately causing cell death. This phenotype then translated to the bulk observation of reduced phage yield, leading us to reassess the results in the previous studies of aminoglycoside antibiotics inhibiting phage propagation ^19,26,39^. We suspect that this may arise from a similar mechanism involving abnormal injection and phage-induced membrane permeabilization that leads to subsequent cell death.

To test this idea, we first examined the phage DNA injection and bacterial physiology by live-cell microscopy. To minimize the effect of phage replication, we used an *E. coli* strain lysogenic for λ (LZ613), in which the prophage carries a kanamycin resistance gene. Similar to our observations with daunorubicin treatment (**Fig. 3h** and **Extended Data Fig. 5a**), live-cell microscopy showed that phage λ failed to complete DNA injection in the presence of kanamycin, an aminoglycoside; λ gDNA did not show any obvious dynamics in the cell (**Fig. 5a, Extended Data Fig. 7a**). This disruption of the injection led to growth arrest without productive phage lysis (**Fig. 5a, Extended Data Fig. 7a**). Similar results were observed in the reporter strain LZ2001, which carries a plasmid conferring kanamycin resistance (**Extended Data Fig. 7b**). The physiological and morphological changes induced by phage-kanamycin combination closely resemble those observed when phages are combined with daunorubicin (**Extended Data Fig. 7c,d**), suggesting a similar cell envelope damage and shared mechanism of action. It is important to note that such killing by the combination of aminoglycosides and phage showed a phenotype distinct from being killed by the antibiotics or the phage alone. Instead of cell lysis or cumulative growth inhibition, phage-kanamycin treatment caused immediate growth arrest, with no elongation, division, or productive phage lysis (**Fig. 5a, Extended Data Fig. 8a,** and **movie S4**).

**Figure 5.**
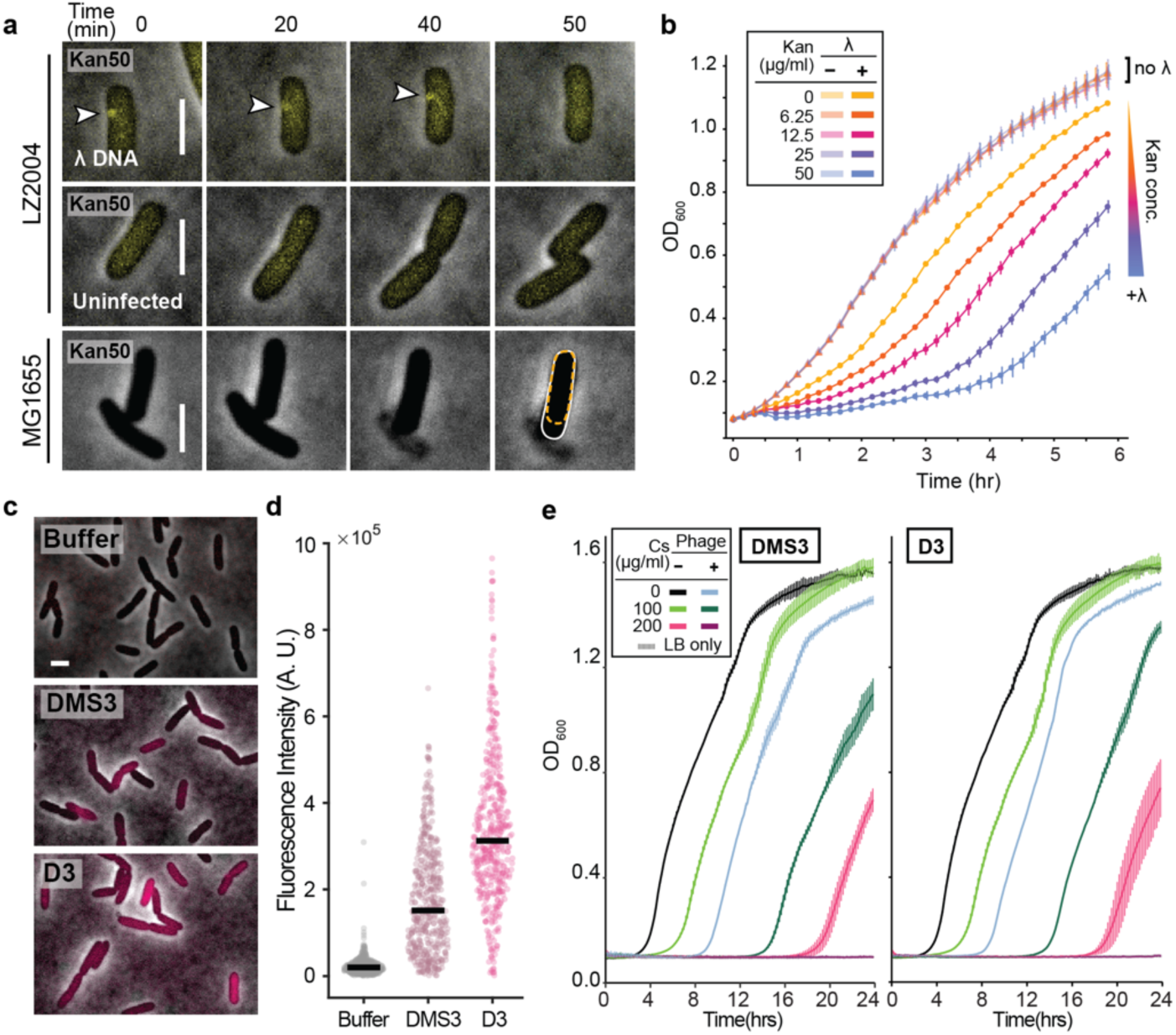
| Synergetic killing of the cell by the combination of phage and antibiotics. **a)** Representative snapshots from a time-lapse movie showing a lambda-infected cell (LZ2004), indicated by successful DNA injection (the yellow focus with an arrowhead pointed to). The infected cell (top) exhibited growth arrest, while the uninfected cell (middle) in the same experiment continued to grow and divide normally. Wildtype MG1655 cells served as the control, showing that non-resistant cells were killed when grown with kanamycin (bottom). The cells continued to elongate (orange dashed lines indicate the cell contour at 0 min, while white solid lines indicate 50 min) before exhibiting growth inhibition or lysis, indicative of cumulative proteotoxic stress. All imaging was performed on M9 + 0.4% maltose agarose pads containing kanamycin at 50 μg/mL. See also **Extended Data Fig. 8a and movie S4**. **b)** Growth curve of both kanamycin- and lambda-resistant *E. coli* (LZ613) under the treatment of phage-drug combination treatments at various sub-inhibitory (<50 μg/mL) concentrations of kanamycin. Wildtype phage λLZ613 was added to an MOI of 5, showing a synergistic effect that sensitizes the bacteria to kanamycin. **c)** DMS3- and D3-infected *P. aeruginosa* PAO1 cells showed a significant increase in daunorubicin accumulation at 10 minutes post-infection in LB medium. **d)** Quantification of daunorubicin signal in individual cells shown in **d**). Number of cells in the quantification: buffer, 475; DMS3, 450; D3, 475. **e)** Growth curve of *P. aeruginosa* cultures treated with phages (D3 and DMS3) and the antibiotic cycloserine at various concentrations. The combination of phage and antibiotic treatment showed a synergistic effect. Scale bars = 2 μm.

Having confirmed that phage infections resensitize kanamycin-resistant *E. coli* and induce growth arrest at the single-cell level, we next tested the effect at the population level by exposing antibiotic-resistant cells (LZ2001) to phage-antibiotic combinations. Supporting the microscopy observation, the inhibition of cell growth under the combined treatment of phage and antibiotics is distinct from the effects of kanamycin or phage alone, where neither cumulative proteotoxic stress (decreased growth over time) nor lysis (drop of OD) was observed, but rather a flat growth curve indicative of an immediate growth inhibition (**Extended Data Fig. 8b**). A similar observation has been made when using a virulent phage, T5 (**Extended Data Fig. 8c**). To avoid potential biases caused during phage propagation and lysis, we further evaluated cell viability under the same condition with the lysogenic strain LZ613. Such a lysogenic strain is resistant to both phage and kanamycin. Monitoring the growth curve revealed that a kanamycin concentration up to 50 μg/mL, which alone had no effect on the lysogen growth, became inhibitory when combined with phage treatment (**Fig. 5b, Extended Data Fig. 8d,** and **Supplementary Fig. 3**).

Having established phage-induced permeabilization in *E. coli*, we next asked whether it also applies to other bacterial host-phage pairs, as well as to antibiotic families beyond aminoglycosides. We sought to use daunorubicin as an indicator to first validate the phage-induced permeabilization of *P. aeruginosa* PAO1 cells by two *Pseudomonas* siphophages, D3 and DMS3. We found that both D3- and DMS3-infected cells showed increased cellular accumulation, similar to that of lambda (**Fig. 5c, d, Supplementary Fig. 4**). To determine if the synergy extends to other antibiotics, we tested the phage in combination with cycloserine (a cell wall synthesis inhibitor) in a liquid growth experiment. We found that the phage-antibiotic combinations caused enhanced growth inhibition compared to either phage alone or cycloserine alone (**Fig. 5e**). Even at a concentration of 200 µg/mL, cycloserine alone did not completely inhibit *P. aeruginosa* growth, whereas the combinations of either DMS3 or D3 at an MOI of 5 with this concentration of cycloserine caused complete growth inhibition (**Fig. 5e**). The enhanced efficacy of phage-cycloserine combinations could also be observed at lower MOIs of 1 and 0.1 (**Supplementary Fig. 5 and 6**). To evaluate whether these results were due to synergistic interactions or were simply due to the phage and antibiotic acting independently, we performed a two-way analysis of variance (ANOVA) as described ^40^. We found that there were statistically significant interactions between DMS3 and cycloserine (**Table S1**) and between D3 and cycloserine (**Table S2**) at the p < 0.0001 level of significance. Similar to what has been observed for lambda with aminoglycosides, the synergistic effects of D3 with cycloserine cause disruption of cell physiology in a way that is distinct from either treatment alone (**Extended Data Fig. 8e**).

## Discussion

In this study, we employed the intrinsic fluorescence of several DNA-intercalating compounds and developed a fluorescence microscopy-based method to assay the uptake and accumulation of small molecules along with monitoring the phage infection process. We found that the level of phage propagation inhibition is related to the fraction of cells undergoing compound-dependent growth arrest rather than the form of incoming phage gDNA or its direct inhibition by the intercalator, unlike the previously suggested mechanism ^19,39^. In particular, our data suggest that phage infection induces permeabilization of the cell envelope to otherwise excluded compounds (**Fig. 1**), thereby reducing the fraction of infective centers that lead to a productive cycle of phage production.

Such phage-induced cell permeabilization resembles observations that a de-energized IM affects the OM and results in the permeation of certain lipophilic compounds, such as N-phenyl-1-naphthylamine ^41^. These compounds are excluded from Gram-negative bacterial cells not only by the OM permeability barrier but also by active efflux pumps ^42^. Supporting this notion, our data show that the apparent impermeability of *E. coli* grown in the presence of daunorubicin results from active efflux mediated by AcrAB (**Fig. 3a-c**), consistent with previous evidence that AcrAB actively effluxes doxorubicin, a daunorubicin homolog ^36^.

This de-energization and permeability increase is also reminiscent of the rapid collapse of membrane potential observed during the initial stage of phage infection ^4^. It must be realized, however, that under normal infection conditions, membrane depolarization and permeability increases are transient and must be recovered to support phage growth. Our data suggest that when phage injection is coupled with certain compounds, membrane depolarization becomes persistent rather than transient. Tracking of phage gDNA in infected cells suggests that the injection process is disrupted, causing constant leakage (**Fig. 3h**). Ultimately, the phage-compound combination drives cells into a state from which they cannot recover, either from the phage-induced channel or from the loss of the ionic gradient across the IM.

Regarding phage specificity, infection with phage lambda at a high MOI (∼5) induced a strong and ubiquitous increase in cellular permeability. Nearly all cells exhibited growth arrest with increasing intracellular daunorubicin signals (**Fig. 4**). Similarly, a uniform, albeit weaker, increase in daunorubicin signal was observed in T5-infected cells. In contrast, T4 infection failed to produce such a signal, and infected cells instead underwent normal lysis. T7 infection produced some daunorubicin-accumulating cells, but the overall signal was much weaker than that of lambda, and cell lysis remained prevalent. Again, these single-cell observations with different phages aligned well with the degree of reduction in phage yield during liquid infection in the presence of daunorubicin or propidium iodide (**Extended Data Fig. 6**), supporting our model of chemical inhibition of phage propagation via permeabilization.

While the precise mechanism by which these compounds interfere with phage DNA injection remains unclear, the finding that phage infection increases the net influx of the compound highlights a promising therapeutic strategy. Specifically, we demonstrate that phage-induced cell permeabilization can resensitize antibiotic-resistant bacteria to multiple antibiotics tested. The synergistic effect was observed across distinct classes of resistant bacteria (**Fig. 5, Extended Data Fig. 7, 8**), including *E. coli* harboring plasmid- or prophage-mediated resistance, as well as intrinsically more resistant species such as *P. aeruginosa* that rely on efflux pumps.

Notably, although lysogens are immune to superinfection by related phages ^43,44^, the combination of phages and antibiotics can nevertheless effectively control lysogenic bacteria regardless of their immunity (**Fig. 5**). A previous study ^45^ suggested a synergistic elimination of lysogens when combined antibiotic treatment with superinfecting temperate phages, where the authors proposed a model in which the lysogen was eliminated due to phage infection-induced SOS responses. In contrast, we suggest that such an effect could also be due to the same model of overall changes to the cell envelope integrity.

Lysogenic bacteria are widespread in nature and constitute an important component of the gut microbiome ^46^. Administering antibiotics often triggers prophage induction, leading to the expression of prophage-encoded toxins such as Shiga toxin ^47,48^. Our data provide an alternative approach to controlling pathogenic populations without inducing cell lysis, thereby mitigating the risks of prophage induction and endotoxin release. It is however worth noting that in the case of superinfecting lysogenic populations, without productive phage production, the synergistic effect will be diminished after a long period, as the effect is only present at the beginning (**Fig. 5b**). In this case, introducing the phage multiple times may serve as a potential solution (**Extended Data Fig. 8d**).

A frequent focus of phage studies is to use it as a potential therapeutic strategy to address the urgent issue of multidrug-resistant bacteria ^49,50^. In particular, several works have focused on evolutionary trade-offs that resensitize antibiotic-resistant pathogens, making them susceptible again to antibiotics ^51,52^. Our study, in contrast, demonstrates that phage entry itself is a direct sensitizing step that lowers bacterial resistance to antibiotics, independent of viral replication. This finding could potentially expand our arsenal against pathogenic bacterial populations.

## Materials and methods

The bacterial strains and phages used in this study are listed in **Tables S3** and **S4**, respectively.

### Thermal induction of phage λ from lysogens

Overnight cultures of *E. coli* lambda lysogen (harboring the *cI857* thermolabile allele of the lambda repressor) were grown in LB Lennox supplemented with 10 mM MgSO_4_ (LBM) at 30°C and 225 rpm. The overday culture was diluted 1:100 into 500 mL fresh LBM in a 4 L Erlenmeyer flask and incubated at 190 rpm to an OD_600_ of 0.4. The culture was thermally induced at 42°C in the water bath at 190 rpm for 15 min. Then, the culture was transferred to 37°C at 190 rpm until OD_600_ ∼ 0.1. The crude lysate was treated twice with 2% chloroform, with each treatment followed by centrifugation at 9,000 rpm (17,568 × g) for 20 min using a Fiberlite F9-6×1000 LEX rotor in a LYNX 6000 centrifuge to pellet cell debris, yielding a clarified crude lysate.

### Preparation of Pseudomonas phages

Overnight cultures of *P. aeruginosa* strain PAO1 were grown in MOPS EZ rich defined medium (Teknova M2105) supplemented with 0.4% glucose, 1 mM of calcium chloride and 10% LB Lennox (EZGluCaLB). For the overday culture, the overnight culture was diluted 1:100 into 50 mL EZGluCaLB in a 500 mL baffled flask. The culture was grown to an OD_600_ of 0.4 before adding phage stocks at a multiplicity of infection (MOI) of ∼0.5. The culture was incubated without shaking for 5 min and allowed to undergo mild shaking at 200 rpm for an additional 4 hours before harvesting. The crude lysate was centrifuged at 20,000 ×g for 20 min twice, followed by filtration through a 0.22 μm membrane filter.

### General procedure of phage enrichment and purification

DNase I (Sigma-Aldrich DN25) and RNase A (Sigma-Aldrich R5500) were added to the clarified lysate at a final concentration of 1 mg/L, and the mixture was incubated at room temperature with gentle shaking (Thermo Scientific rotator model 2309) for 1 hour. Depending on the volume, crude lysates were precipitated with 10% PEG-8000 in 1M NaCl, or enriched through differential sedimentation. Smaller-volume samples were enriched by centrifugation at 20,000 rpm (51,428 × g) for 3 hours using a Fiberlite F20-12×50 LEX rotor in a LYNX 6000 centrifuge, whereas larger-volume samples were centrifuged using a Fiberlite F9-6×1000 LEX rotor at 9,000 rpm (17,568 × g) for 15 hours. The supernatant was discarded and the phage pellet was recovered by soaking in a small volume (∼1-5 mL) of gelatin-free SM buffer for 6 hours at 4°C. The efficiency of differential sedimentation was routinely verified by titering the plaque-forming units in supernatant in comparison to the phage pellet resuspension. The resuspended phage solution was further purified by centrifuging at 18,000 ×g for 10 minutes and subjected to isopycnic centrifugation in CsCl solution. The lysate was mixed with a 1.5 g/mL CsCl stock solution in gelatin-free SM buffer in a Beckman 13.2 mL UltraClear tube to achieve a final density of ∼1.45 g/mL. The tube was subjected to ultracentrifugation at 35,000 rpm with an SW41Ti rotor for 48 hours. The phage band was extracted using a beveled 18G needle and dialyzed against 1 liter of SM buffer in three rounds for 6 hours, overnight (∼12-16 hours), and an additional 6 hours.

### Microscopy of phage infection in the presence of small-molecule compounds

Overnight cultures of MG1655 were routinely grown in LB medium at 37°C and 265 rpm. The next day, the culture was diluted 1:100 into 5 mL of fresh LB medium in a baffled flask and grown to an OD_600_ of 0.1. Small molecules were then added, and the culture was allowed to continue growing to an OD_600_ of 0.4. For the infection, 50 µL of cells (2 × 10^8^ CFU/mL, CFU for *<u>c</u>*olony *f*orming *<u>u</u>*nits) were mixed with 20 µL of a phage dilution (2.5 × 10^9^ PFU/mL, PFU for *<u>p</u>*laque *f*orming *<u>u</u>*nits) to achieve an MOI of 5. The mixture was incubated in a 30°C water bath for 10 minutes before imaging.

For experiments with *P. aeruginosa*, overnight cultures of PAO1 were routinely grown in LB medium at 37°C and 265 rpm. The next day, the culture was diluted 1:100 into 10 mL of fresh LB medium in a baffled flask and grown to an OD_600_ of 0.1. Daunorubicin was then added to a concentration of 15 µM, and the culture was allowed to continue growing to an OD_600_ of 0.4. For the infection, 50 µL of cells (3 × 10^8^ CFU/mL) were mixed with 5 µL of a phage dilution (1.5 × 10^10^ PFU/mL) to achieve an MOI of 5. The phage dilution was prepared in phosphate-buffered saline (1 × PBS). The mixture was incubated in a 37 °C water bath for 10 minutes before imaging.

### Phage propagation in liquid culture with small-molecule compounds

The inhibitory function of the small-molecule compounds on phage propagation was tested based on the previously reported method ^19^. MG1655 cultures were grown overnight at 37 °C in LB medium. The next day, the overnight culture was diluted 1:100 into 5 mL fresh LB medium, with or without the addition of 15 µM Daunorubicin, 40 µM propidium iodide. The cultures were incubated at 37 °C and 265 rpm for 1 hour. Following this, phage (∼4 x 10^9^ PFU) was added at a multiplicity of infection of ∼10. The cultures were then incubated under the same conditions for an additional 6 hours. The supernatant was collected by centrifugation at 15,000 rpm for 2 minutes, and the phage titer was determined by spotting serial dilutions onto a lawn of MG1655 on LB agar plates. Three biological replicates were tested for each phage.

### Constructing manY/Z reporter strains

To visualize that the phage DNA is being ejected into the host cytoplasm in the absence of the IM receptor (ManYZ), we introduced the TetR-mNeongreen reporter plasmid into the *manY* and *manZ* single-gene knockout strains. The parental *manY* and *manZ* strains (RY33002 and RY33003, see **Table S3** for more details) were originally constructed by transducing the *manY*/*manZ*::*kan^R^* construct from the Keio collection strains ^53^ into the MG1655 background. The reporter plasmid pZS*34-*P*_FtsK*i*_*-tetR-mNeonGreen* ^25^ was then transformed into chemically competent cells using standard TSS transformation.

### Sensitize PL15 strain for phage T5 infection

The parental strain of the sticky flagella strain PL15 is RP437 (CGSC #12122), which harbors a mutation of *fhuA31* that abolishes the ability for T1 and T5 to infect the cell. To revert *fhuA31* to the wild-type state to sensitize the strain, we did P1 transduction (see below) to replace the adjacent upstream gene, *mrcB*, with the *mrcB*::*kan^R^*(transduced from the Keio collection single-gene knockout strain JW0145 ^53^. Through co-transduction, the wild-type fhuA gene is co-transduced with *mrcB*::*kan^R^* and thereby reverts fhuA31 to the wild-type state. The outcome strain (LZ3543) was phenotypically and sequence verified.

### P1 transduction

To generate a P1*vir* lysate containing transducing particles, overnight cultures of donor strains (e.g., Keio collection knockouts) were diluted 1:200 into 3 mL LB supplemented with 10 mM MgSO₄ and 5 mM CaCl₂ and grown at 37 °C with shaking. Once cultures reached early log phase (OD_600_ ∼0.2-0.25), ∼5×10^7^ PFU of P1*vir* were added to initiate infection. After 2–3 hours of incubation, a clearance of the culture should be evident and the lysates were clarified and collected by chloroform treatment and centrifugation, and stored at 4 °C for use in the transduction steps.

Recipient cells were grown overnight in LB and prepared for direct use by adding 10 mM MgSO₄ and 5 mM CaCl₂. Infections were set up by mixing 100 μL of recipient cells with 50 μL of series dilutions of the transducing lysate, followed by incubation at 37 °C for 30 minutes without shaking. Sodium citrate was then added to a final concentration of 100 mM to chelate calcium and prevent further phage infection. After an additional 60 minutes of incubation, mixtures were centrifuged, resuspended in LB supplemented with 100 mM sodium citrate, and plated on selective plates containing 20 mM sodium citrate. Colonies were re-streaked at least twice on the same sodium citrate selective plates to eliminate residual P1*vir* and isolate purified transductants.

### Live-cell imaging in suspension

Live cell infection movies of T-phages were carried out in cell suspension using the sticky flagella strain PL15 *motAB* ^54^. In brief, the strain harbors a sticky *fliC* allele (*fliC*^sticky^) that allows flagella to bind to the glass surface for immobilization. Additionally, the strain has mutations in the stator of the flagellar motor (*motAB*), which abolishes the spinning of the bacteria. To obtain an optimal nutrient condition for long-term observation of phage infection, glass-bottom Petri dishes were used in this experiment (CellVis D35-10-1-N) to provide a sufficient reservoir to allow cell growth.

In a typical experiment, overday culture of PL15 was grown in a Falcon snap cap culture tube to mid-log phase with OD_600_ ∼ 0.5. To adhere the cell to the glass surface, 200 μL of the overday culture was applied to the center of the 35 mm petri dish where the glass coverslip is located. The culture was spread evenly across the glass surface by pipetting. The petri dish was then left undisturbed for 15 min at room temperature, and the unbound cells in the culture were discarded. This was followed by three rounds of washes with 2 mL LB each round. After the addition of the third batch of LB, the sample was left static for another 15 min prior to the imaging. If the experiment requires phage infections or the supplementation of certain antibiotics/compounds, the sample will be treated with one additional buffer exchange to replace the 2 mL LB with the desired buffer or phage/drug mixture. The samples were placed into a custom heat chamber (Okolab stage top incubator) set to 30 °C and placed under the microscope and imaged under phase-contrast and mCherry (10 ms exposure) every 5 min for 2 hours.

For a typical infection, 10 µL of 2×10^10^ PFU/mL phage dilution was added to the final suspension medium, which gave a final MOI of ∼2. Either the phage dilution or the volume to be added to the buffer can be adjusted if a higher MOI is needed. In order to rule out the possibility of cell killing by ‘lysis-from-without’, a negative control with only LB and phage at the same concentration was used to make sure normal cell lysis could be achieved at such a high MOI.

### General microscopic imaging settings

Microscopy for qualitative and quantitative purposes were performed on either a Nikon Eclipse Ti inverted epifluorescence microscope or a Nikon Eclipse Ti2 inverted microscope. For the Nikon Eclipse Ti, a 100 × objective (Plan Fluor, NA 1.40, oil immersion) was used with a 2.5× TV relay lens. Imaging was performed using either a mercury lamp or a SOLAR Engine III LED as the light source and acquired using a cooled EMCCD (electron-multiplying charge-coupled device) camera (iXon3 897, Andor, Belfast, United Kingdom). For the Nikon Eclipse Ti2, a 100 × objective (Plan Fluor, NA 1.45, oil immersion) was used. Fluorescence was excited using an X-Cite XYLIS LED illumination system. Depending on the desired field of view, images were sometimes acquired with a 1.5× microscope built-in magnification lens. Images on Nikon Eclipse Ti2 microscope were captured using either a Princeton Instrument ProEM-HS EMCCD camera or a Hamamatsu ORCA-Fusion sCMOS camera (Hamamatsu Photonics C14440-20UP).

### Image processing and analysis

Microscope images were exported from Nikon Elements AR software as raw TIFF files. Cells were segmented from phase-contrast images using either a custom U-Net model^55^ or Omnipose (https://github.com/kevinjohncutler/omnipose) ^56^. Output files were saved and analyzed using custom MATLAB scripts.

### Use of artificial intelligence (AI) and large language models (LLM)

LLMs (ChatGPT and Google Gemini) were used solely for copy-editing and proofreading of human-generated text and human-written MATLAB scripts. Both the AI-edited code and the original code (retained as comments) are provided. No other use of LLMs or generative AI was involved in this study.

### Microplate-based quantification of phage-antibiotic synergy at the population level

To assess the combinatorial effects of phages and antibiotics on bacterial growth, a 96-well microtiter plate assay was conducted using the Tecan Spark plate reader. *Pseudomonas aeruginosa* strain PAO1 wildtype was grown overnight at 37 °C in LB medium with shaking at 265 rpm. The next day, a 1:100 dilution of the overnight culture was inoculated into fresh LB medium and grown to mid-exponential phase (OD_600_ ∼ 0.4), corresponding to approximately 3 × 10^8^ CFU/mL. During this time, working solutions of antibiotics were prepared in LB at 2× the desired final concentrations, and high-titer phage stocks were diluted in 1× PBS to yield the desired MOI. Control dilutions using buffer without phage were also prepared.

For each experimental condition, 75 µL of antibiotic solution and 75 µL of inoculum (overday culture diluted to 1 × 10^6^ CFU/mL bacteria, with or without phage) were combined in triplicate wells, giving the final inoculation density in each well at 5 × 10^5^ CFU/mL. This inoculum density was chosen based on established best practices for antimicrobial susceptibility testing, as detailed in methods published by the Clinical and Laboratory Standards Institute ^57^. Controls included wells with bacteria but no phage or antibiotic (growth control), and wells with LB only (sterility control). After loading, the plate was placed into the Tecan Spark plate reader and a kinetic measurement protocol was initiated to collect absorbance data at 600 nm every 10 minutes for 24 hours (144 cycles total), with temperature maintained at 37 °C and double-orbital shaking at 90 rpm.

The combinatorial effects of coliphages and aminoglycosides were assessed using a similar workflow. Briefly, the λLZ613 lysogen strain was grown overnight at 37 °C in LB supplemented with kanamycin. The following day, the culture was inoculated 1:100 into 10 mL of fresh LB and grown to mid-log phase (OD_600_ ∼ 0.4). The culture was then back-diluted in LB to approximately 1 × 10⁸ CFU/mL. Cells were mixed with phage or buffer to achieve the desired MOI and immediately transferred into a 96-well plate pre-loaded with 2x antibiotics. The growth kinetics were monitored for 24 hours at 30 °C using the Tecan Spark plate reader with continuous shaking.

To evaluate the effects of daunorubicin on the growth of *E. coli* MG1655, an overnight culture was diluted 1:100 into fresh LB medium in the flask and grown to early-log phase OD_600_ ≈ 0.1, followed by the addition of either the daunorubicin stock to reach the working concentration or an equivalent volume of buffer control. The mixture was then transferred to a 96-well plate. To disrupt the OM function, EDTA or buffer control was directly added to the wells and mixed with the cells. The growth kinetics were monitored for 6 hours at 37 °C using the Tecan Spark plate reader with continuous shaking.

## Acknowledgements

We thank KC Huang, Pushkar Lele, Jason Gill, and the Center for Phage Technology for sharing strains. We are grateful to Alan Davidson, Karen Maxwell, and Annie Si Cong Li for sending *Pseudomonas* phages and thoughtful discussion on early versions of the results. We thank Paul Straight and all members of the Zeng lab for their insights. We are grateful to Ido Golding for commenting on the earlier version of the manuscript. This work was in part supported by NSF under Grant No. MCB-2013762. HRL is supported by the NSF Graduate Research Fellowship Program under Grant No. 2139772. LT was supported by the NSF Research Experiences for Undergraduates (REU) program under Grant No. 1949893. Any opinions, findings, and conclusions or recommendations expressed in this material are those of the authors and do not necessarily reflect the views of the National Science Foundation.

## Data and materials availability

Data for individual replicates of fluorescence quantifications are available in Supplementary Figures 1 and 3-6, and in the Source Data. Raw image files and custom scripts for image analysis are available in the Zenodo repository associated with the manuscript. Statistical analyses are summarized in **Tables S1 and S2**. Bacterial strains and bacteriophages used in this study are listed in **Tables S3 and S4** and are available upon request.

## Author contributions

Conceptualization: Z.Y., R.Y. and L.Z. Investigation: Z.Y., C.W., H.L., I.M., L.T., R.Y. and L.Z. Funding acquisition: L.Z. Supervision: L.Z. Writing – original draft: Z.Y. and L.Z. Writing – review and editing: Z.Y., C.W., H.L., R.Y. and L.Z.

## Extended Data Figures

**Extended Data Fig. 1.**
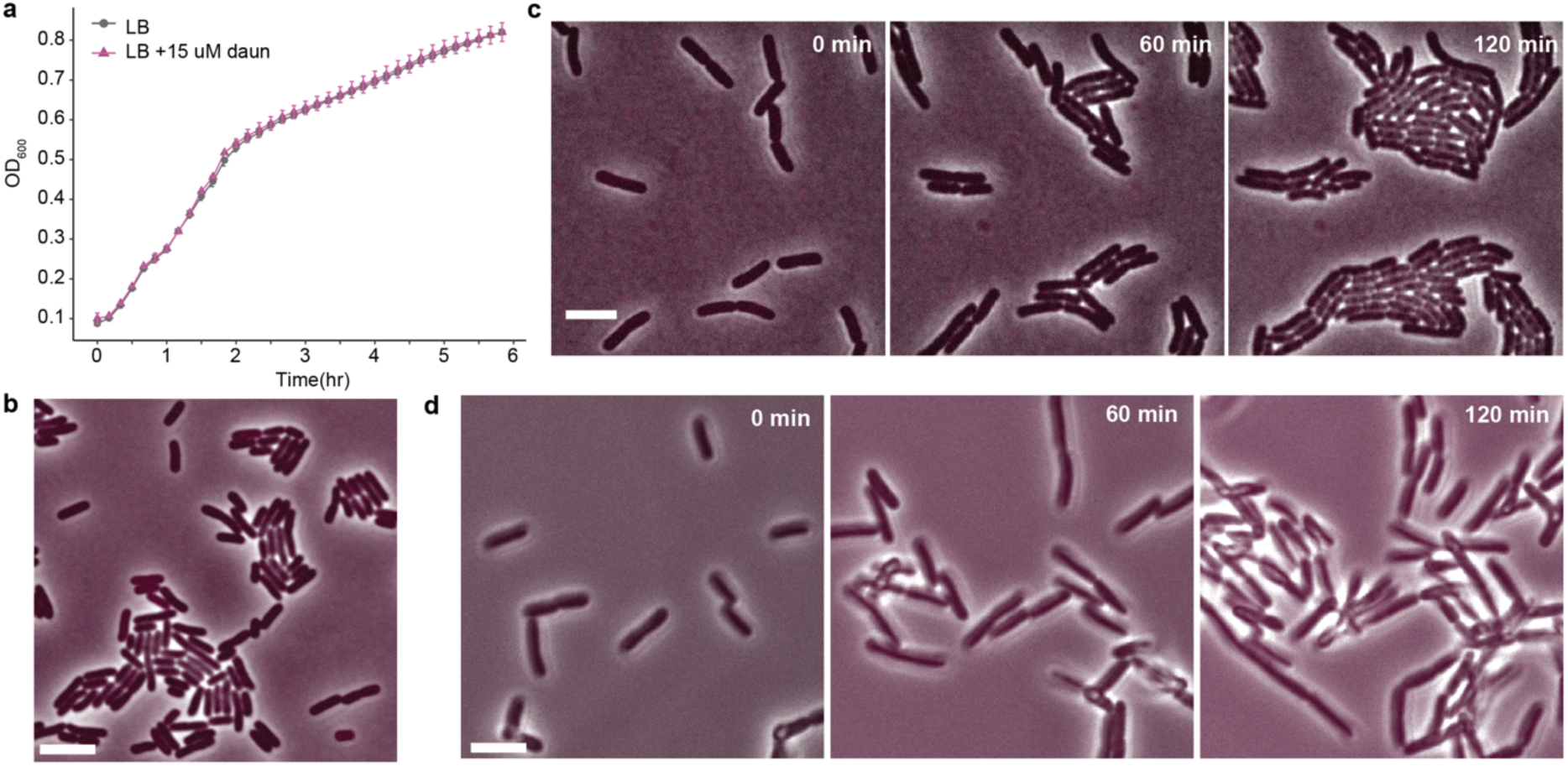
| Cells grow normally in the presence of daunorubicin. **a)** The presence of daunorubicin does not affect *E. coli* MG1655 growth. **b)** An exponentially growing culture of MG1655 in the presence of daunorubicin showed no intracellular accumulation of daunorubicin. **c)** Representative snapshots from a time-lapse movie of MG1655 growing and dividing on an LB + daunorubicin (15 µM) agarose pad. **d)** Representative snapshots from a time-lapse movie of PL15 growing and dividing in suspension on LB + daunorubicin (15 µM), adhered to the surface of a glass-bottom dish. Scale bars = 5 µm.

**Extended Data Fig. 2.**
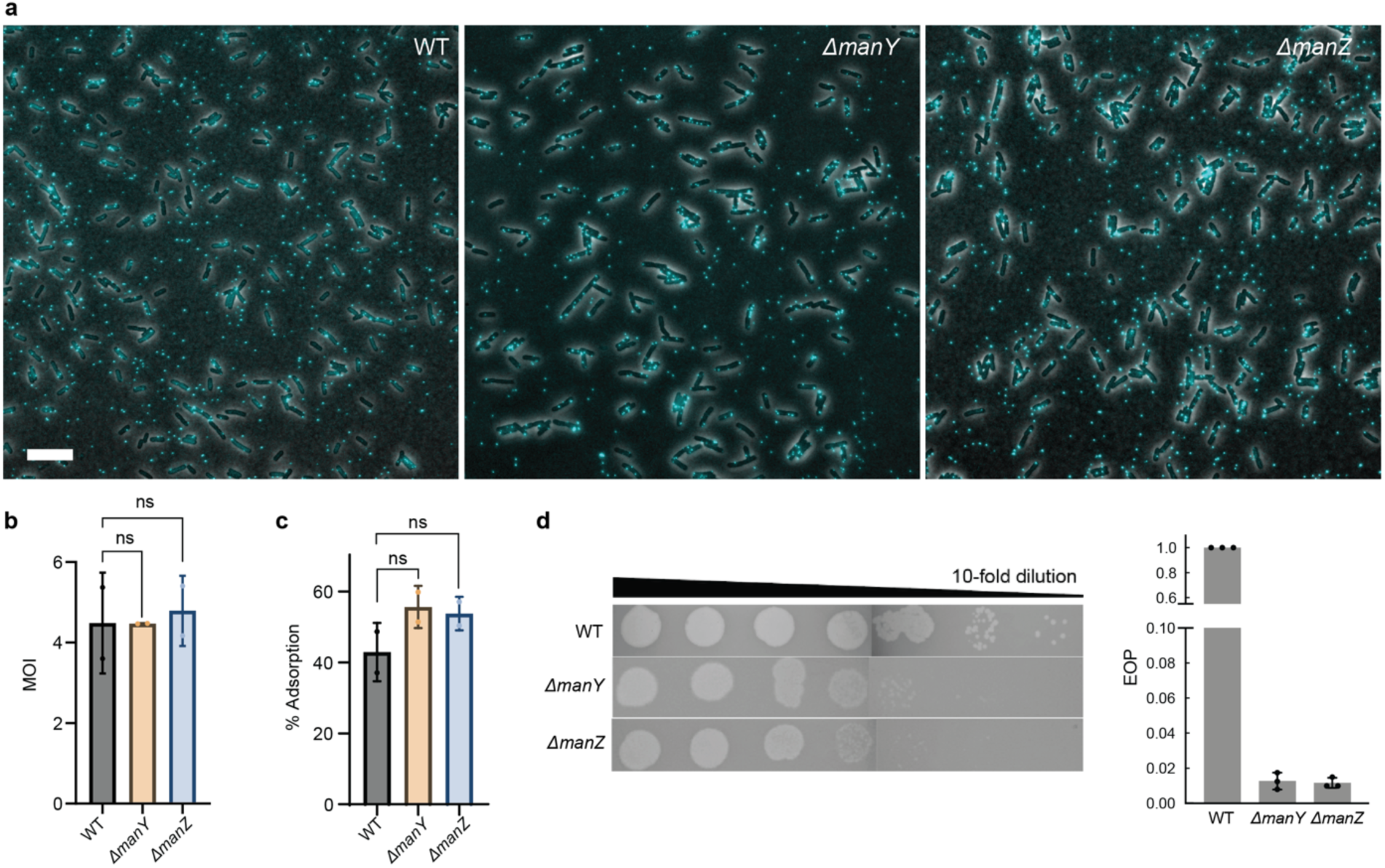
| Plaquing efficiency comparison between wildtype and *ΔmanY/ΔmanZ* mutants. **a)** Phage λLZ2002 adsorbs similarly to the *ΔmanY*/*ΔmanZ* mutants as compared to the wildtype. Individual phage particles are visualized as cyan dots either on the cell surface or as free virions in the background. **b)** Quantification of the infection MOI for **a)**, which represents the total phage amount over cells counted from the microscope images. **c)** Quantification of phage λLZ2002 adsorption on the *ΔmanY/Z* mutant strains compared to the wildtype. Statistical significance was determined via ordinary one-way ANOVA, where two biological replicates were treated as repeated measures, and the mean value of each group was compared to the mean of the control group (wildtype). P-values calculated as 0.5602 and 0.1923 for *ΔmanY* and *ΔmanZ*, respectively. **d)** The *ΔmanY* and *ΔmanZ* mutant strains (LZ3526 and LZ3527) used in this study have a plating efficiency for λLZ613 at ∼50-fold decrease from the WT (LZ2001). These two mutants produce slightly smaller and fuzzier plaques.

**Extended Data Fig. 3.**
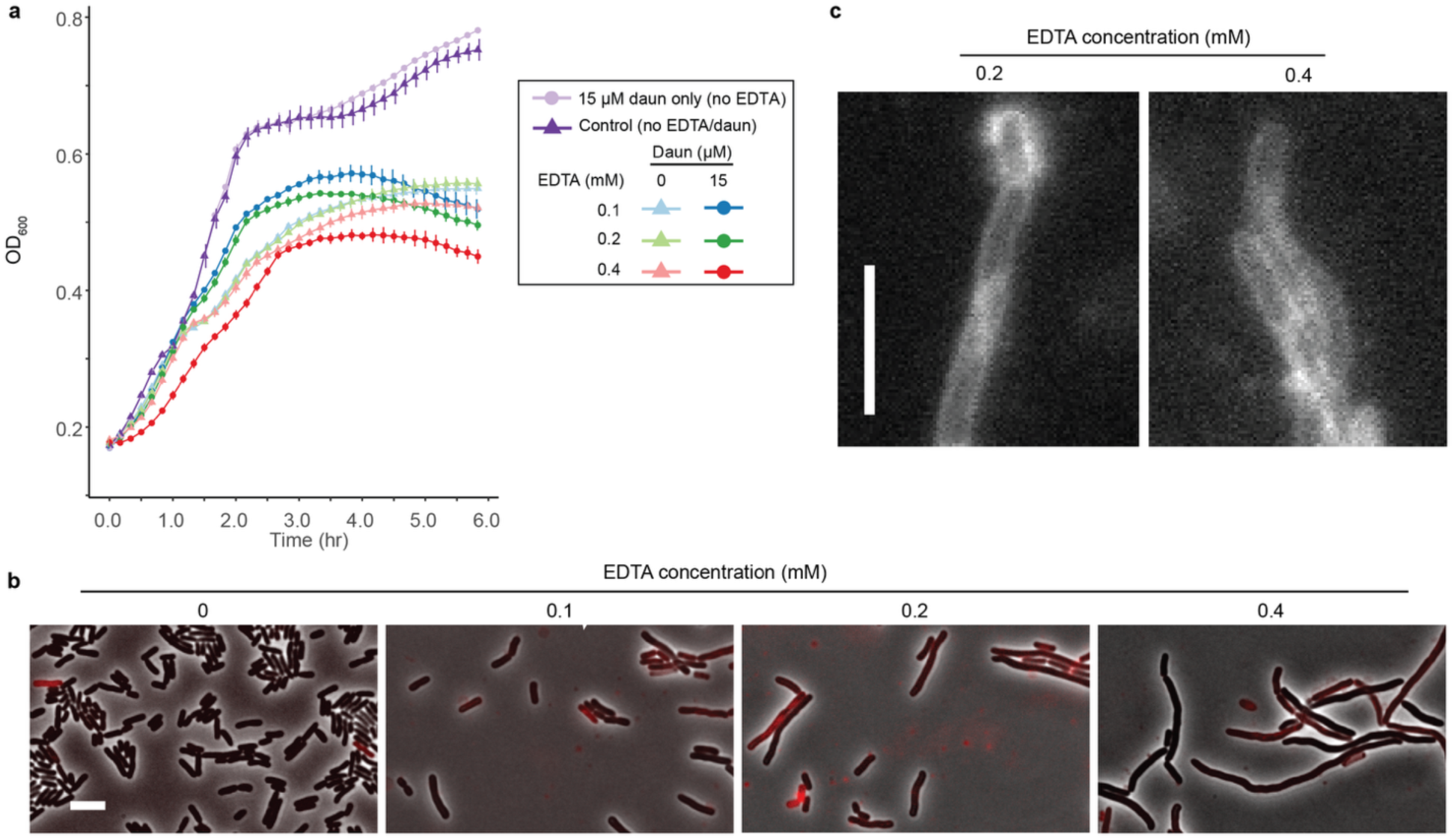
| The effect of EDTA on changing daunorubicin permeability. **a)** Plate reader measurement of cell growth in the presence of EDTA and daunorubicin. **b)** Although the addition of EDTA seems to inhibit cell growth when combined with 15 µM daunorubicin, cells under this combination of treatment show different patterns of intracellular daunorubicin signals compared to phage infections. With the increase of concentrations of EDTA, cells undergo more severe growth defects and show filamentation. Compromised cells then show increased daunorubicin signals. **c)** Zoomed-in view of the daunorubicin signal channel showing representative filamented cells in **b**). These cells with growth defects exhibit only weak daunorubicin signals and show ring-like accumulation patterns. Scale bars = 5 µm.

**Extended Data Fig. 4.**
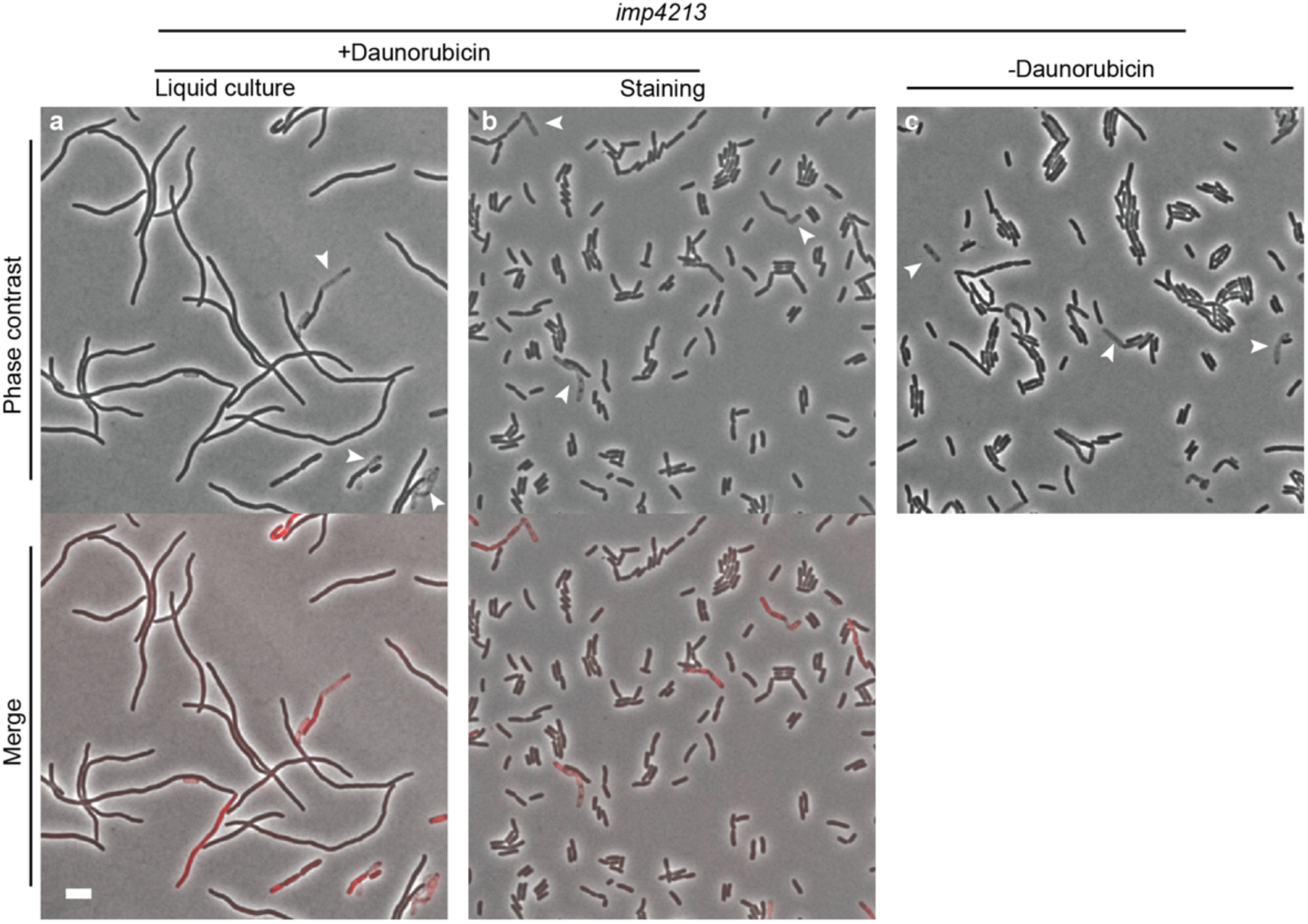
| The effect of an impaired outer membrane on changing the cell permeability. *imp4213* (KC427) cells grown and imaged under different conditions may show autolysis (phase light cells indicated by white arrowheads), filamentation, and staining by the daunorubicin, specifically: **a)** Cells grown in LB+15 µM daunorubicin for 1.5 hours showed filamentation without high-level accumulation of daunorubicin in the cells. Red cells may correspond to the lysed or compromised cells, as suggested by the correlation between red cells and the phase light cells. **b)** Cells grown in LB and stained with 15 µM daunorubicin for 30 min showed only a limited amount of daunorubicin signal in the majority of the cells. Similar to the liquid culture condition shown in **a)**, the red cells may be correlated with compromised cells. **c)** Autolysis is evident under normal growth conditions without daunorubicin, indicating a general impaired cell integrity. The lookup tables (LUTs) for the bright-field channel in this figure were specifically adjusted to appear brighter (compared to other images in this study) for better visualization of the phase-light cells. Scale bar denotes 5 µm.

**Extended Data Fig. 5.**
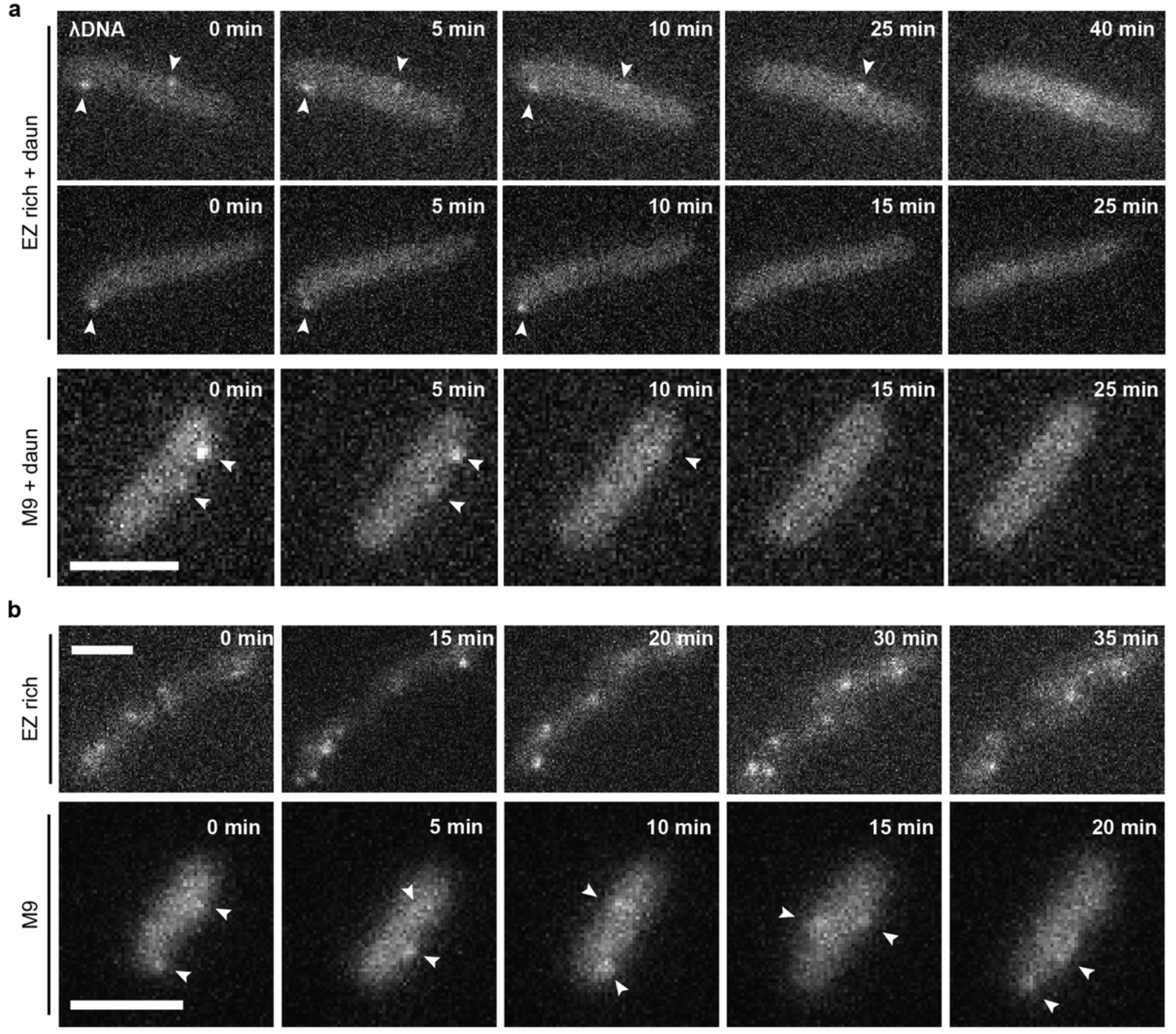
| Phage gDNA penetration does not complete in the presence of daunorubicin. **a)** Snapshots from time-lapse movies capturing three representative cells in EZ-rich defined media supplemented with 15 µM daunorubicin and infected with phage λLZ2002. Fluorescent puncta, indicated by white arrowheads, represent internalized lambda DNA in the cytoplasm. The fluorescent reporter specifically binds to the *tetO* array at the right end of the phage λLZ2002 gDNA, which enters the cytoplasm first. Reporter-bound phage gDNA appeared static for a prolonged period (approximately 10–20 min) before later disappearing. These movies suggest that gDNA penetration occurred in the presence of daunorubicin but was likely incomplete, leaving the right end of the phage gDNA at the site of adsorption and injection. **b)** Representative time-lapse movie showing normal phage infection without daunorubicin. Fully internalized phage gDNA exhibited dynamic movement inside the cell. Scale bars = 2 µm.

**Extended Data Fig. 6.**
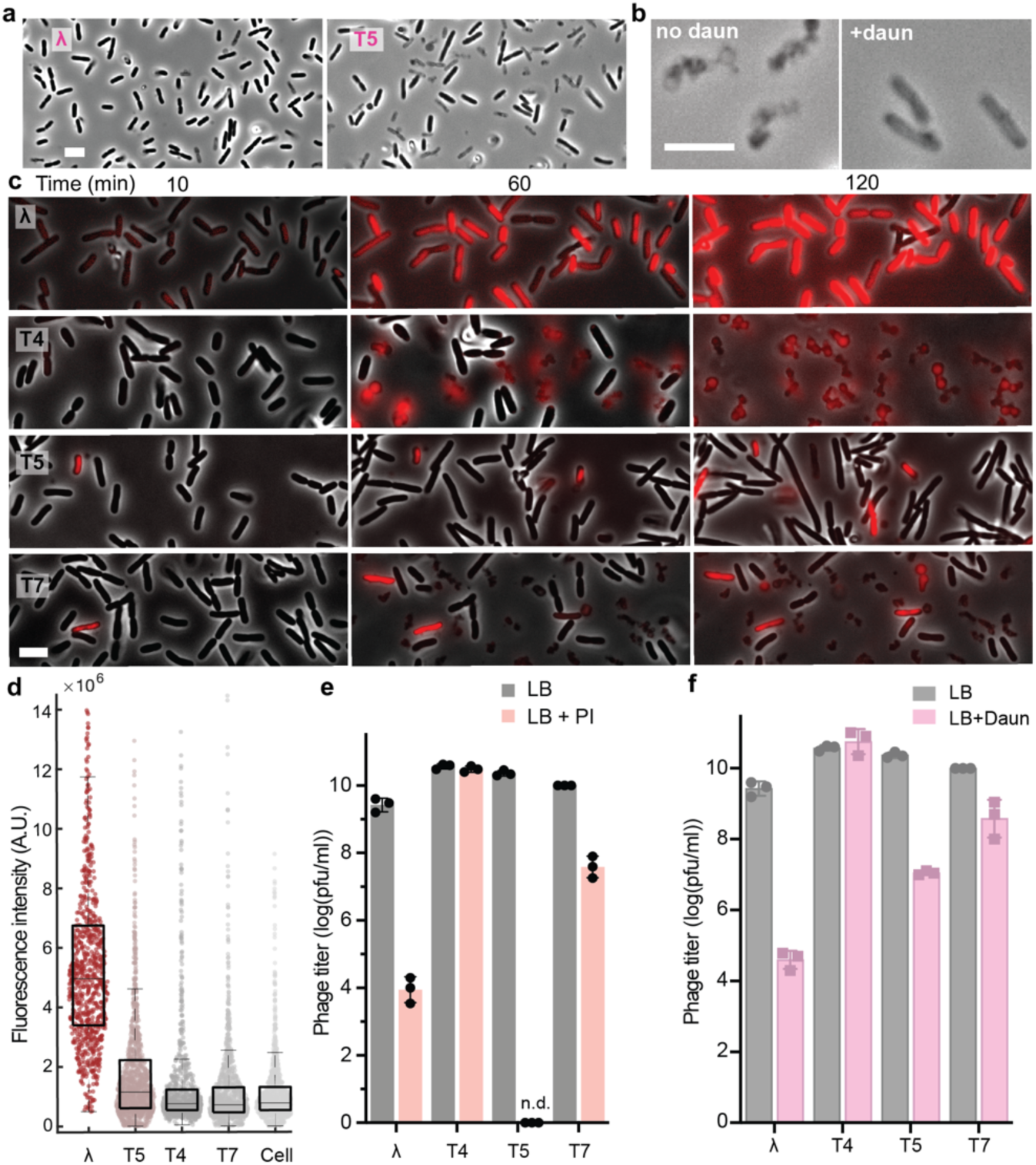
| T-phage-induced daunorubicin or propidium iodide accumulation correlated with less productive phage production in a liquid propagation experiment. **a)** Unlike λ, the T5-daunorubicin combination showed many phase-light ghost cells. Snapshots of the phase-contrast channel were taken at 120 min post-infection from the same dataset as Fig. 4a. **b)** Phage T5 infection in combination with daunorubicin induces a unique phase-light ghost phenotype, distinct from lysis during normal infection without daunorubicin. **c)** Representative snapshots of PL15 cells over time with propidium iodide (PI) accumulation when infected by λLZ613, T4, T5 and T7 at high MOIs (>2). **d)** Quantification of intracellular signals of propidium iodide at 10 min post-infection with an MOI of 5. **e, f**) Reduction of phage-productive infection reflected in the drop of EOP when propidium iodide (**e**) or daunorubicin (**f**) was added to the infection mixture. Scale bar = 5 µm. n.d. = not detected.

**Extended Data Fig. 7.**
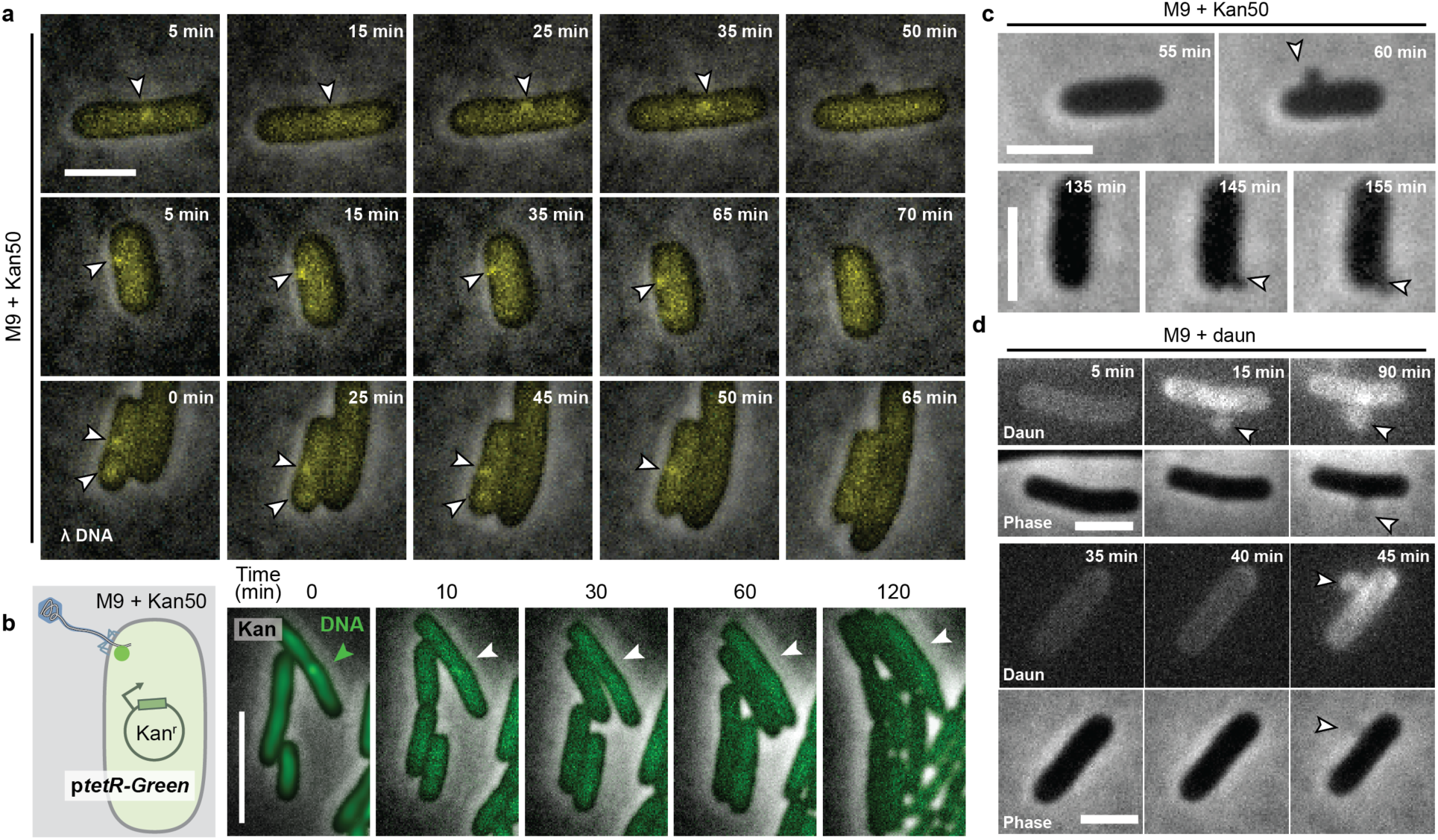
| Evidence that phage λ infection in the presence of kanamycin leads to partial DNA injection, similar to that observed with daunorubicin treatment. **a)** Four representative cells lysogenic for lambda (LZ2004) superinfected by the reporter phage λLZ2002. The lysogen LZ2004 harbors the λLZ613 prophage, which confers kanamycin resistance. λLZ2002 gDNA penetrating into the cytoplasm appears as fluorescent yellow puncta (indicated by arrowheads). In these examples, the DNA foci remained at a fixed position for an extended period (∼1 hr) before eventually fading away, indicative of incomplete gDNA penetration, similar to what was observed with daunorubicin treatment (**Extended Data** Fig. 5a). **b)** Same treatment as in **a**) and Fig. 5a, except the cells carried kanamycin resistance from a plasmid rather than the prophage. Similar results were observed: uninfected cells elongated and divided, while phage-infected cells (indicated by an arrowhead) showed growth arrest, with the DNA foci remaining static and later disappearing. **c)** Phase-contrast images of cells under the same conditions as in **a**), showing damaged cell envelope structures (blebs, indicated by arrowheads of phase contrast images) that likely led to the loss of cellular content. **d)** Similar to kanamycin-treated cells as shown in **b**), cells infected by lambda in the presence of daunorubicin exhibited similar cell envelope damage and loss of cellular contents, suggesting a similar mechanism of action leading to growth arrest. Fluorescence signals indicate the accumulation of daunorubicin inside the cell, but are also present in the blebs (indicated by arrowheads). All scale bars indicate 2 µm. Also see **movie S4** for more examples.

**Extended Data Fig. 8.**
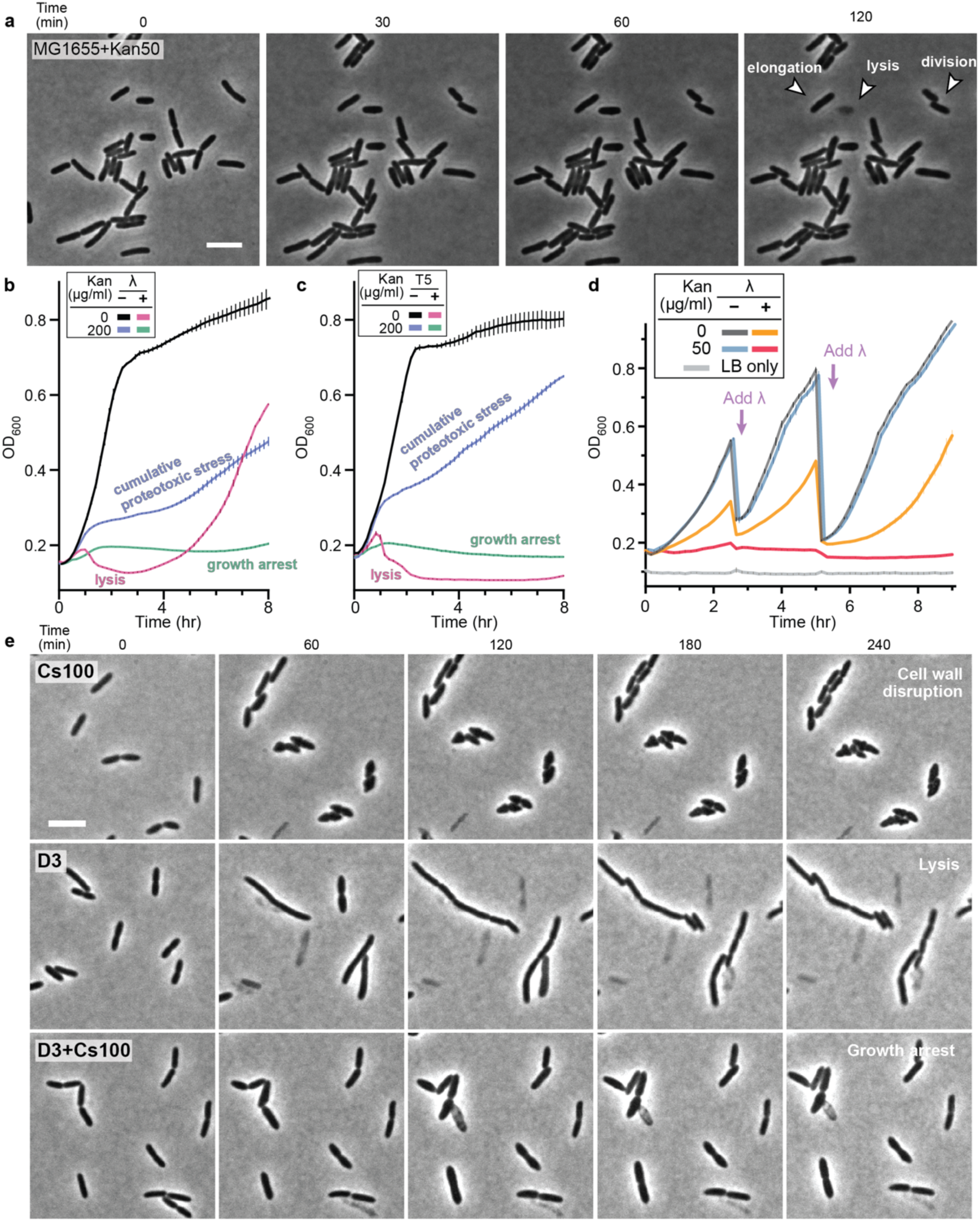
| Synergistic effect of phage-antibiotic combinations under various conditions. **a)** Time-lapse movie showing wild-type *E. coli* (MG1655) grown on a kanamycin (50 µg/mL) agarose pad. Proteotoxic stress was cumulative, and cells were able to grow, elongate, and divide before exhibiting growth inhibition and lysis. **b)** Growth curve of a kanamycin-resistant *E. coli* (LZ2001) under the treatment of lambda-kanamycin combination treatments. Distinct growth profiles were observed for phage treatment alone (lysis; drop of OD), kanamycin alone (accumulated inhibition of growth over time), or the combined treatment (growth arrest). **c)** Growth curve of a kanamycin-resistant *E. coli* (LZ2001) under the treatment of T5-kanamycin combination treatments. Similar to the λ-kanamycin combination **b**), T5-kanmycin combination showed a distinct growth curve as compared to either phage or kanamycin alone. **d)** Growth curve of both kanamycin- and lambda-resistant *E. coli* (LZ613) under the treatment of phage-drug combination treatments at various sub-inhibitory (<50 μg/mL) concentrations of kanamycin. This experiment was conducted in a similar manner to Fig. 5b, but with multiple batches of back-dilutions and phage additions (purple arrows). Each time, wildtype phage λLZ613 was added at an MOI of 5, showing a synergistic effect that sensitizes the bacteria to kanamycin. **e)** Representative snapshots from a time-lapse movie showing the distinct pattern of cell physiology disruption caused by cycloserine alone (abnormal cell morphology), phage D3 alone (lysis), or the combination of the two treatments (growth arrest). Consistent with the observation from the growth curve measurement in Fig. 5e. Scale bars = 5 µm.

