## Supplemental Text, Tables and Figures. for "Lethal permeabilization of host bacteria to small-molecule compounds during phage penetration"

*for*

**List of content:**

Supporting information text, including:

Descriptions for statistical analysis used in Figure 5

Legends for supplementary movies S1 - S4

Supplementary Tables S1 - S4

Supplementary Figures S1 - S6

SI references

**Other supporting materials for this manuscript:**

Supplementary movies S1 - S4

*Supporting information text*

**Statistical analysis used in Fig. 5 to distinguish synergistic from additive effects**

We used two-way analysis of variance (ANOVA) to rigorously assess whether the enhanced inhibition of *P. aeruginosa* growth caused by the phage-antibiotic combinations was due to additive or synergistic effects, as previously described .There were statistically significant interactions between phage DMS3 and cycloserine ($p = 4.62 \times 10^{-21}$) and between phage D3 and cycloserine ($p = 6.28 \times 10^{-15}$) in inhibiting *P. aeruginosa* growth, indicating that both phages exhibited synergy with this antibiotic. For this analysis, growth was quantified using the areas under the growth curves (AUCs). See **Tables S1 and S2** for more information.

**Legends for supplementary movies**

**Movie S1: Phage infection causes daunorubicin accumulation and growth arrest**
Time-lapse imaging of LZ2001 cells infected by λLZ2002 (cyan foci) showed increasing daunorubicin uptake/accumulation (magenta) and growth arrest. A snapshot at time zero is included at the start of the movie, showing the overlay of adsorbed phage particles with injected DNA (yellow foci, reporting the right end of lambda gDNA). Selected frames from the movie are presented as montages in **Fig. 1a**.

**Movie S2: Daunorubicin interfering with the lambda injection process.**
Time-lapse imaging of LZ2001 cells infected by λLZ2002 in the presence of daunorubicin. λLZ2002 gDNA showed no obvious dynamics, suggesting that the injection process is disrupted, leading to persistent depolarization (top panels). Examples showing DNA dynamics observed during λLZ2002 infection of LZ2004 in the absence of daunorubicin were included for comparison (bottom panels). Additional example snapshots are shown in **Extended Data Fig. 5**. See also **Extended Data Fig. 7** for similar observations when phage infection is combined with kanamycin.

**Movie S3: Distinct phenotype of cell permeabilization to daunorubicin/propidium iodide caused by different phages.**
Time-lapse movie showing the comparison of the effect of phage-daunorubicin (left) and phage-propidium combination (right) with different coliphages (λLZ613, T4, T5, and T7).

**Movie S4: Phage-kanamycin combination causes lysogen growth arrest.**
The kanamycin-phage combination causes LZ2004 growth arrest (left). Uninfected cells showed elongation and division due to their kanamycin resistance (left). Phage replication and lysis were included for comparison (middle). Wildtype (MG1655) cells subjected to proteotoxic stress by supplementation with kanamycin (50 μg/mL) were included as a control (right). Together, these examples show that the phage-kanamycin combination kills via a mechanism distinct from phage or kanamycin alone.

**Table S1 | Two-Way ANOVA Results for DMS3 and Cycloserine**

| Source Term | Sum of Squares | Degrees of Freedom | Mean Square | F-statistic | p-Value |
|---|---|---|---|---|---|
| DMS3 MOI | 15663 | 3 | 5221 | 1719.9 | $3.90 \times 10^{-28}$ |
| Cycloserine Concentration | 48131 | 2 | 24066 | 7927.8 | $1.42 \times 10^{-34}$ |
| Interaction Between DMS3 and Cycloserine | 5168.1 | 6 | 861.3 | 283.7 | $4.62 \times 10^{-21}$ |
| Error | 72.9 | 24 | 3.04 | | |
| Total | 69035 | 35 | | | |

**Table S2 | Two-Way ANOVA Results for D3 and Cycloserine**

| Source Term | Sum of Squares | Degrees of Freedom | Mean Square | F-statistic | p-Value |
|---|---|---|---|---|---|
| D3 MOI | 9754.2 | 3 | 3251.4 | 515.3 | $6.51 \times 10^{-22}$ |
| Cycloserine Concentration | 57191 | 2 | 28595 | 4532.3 | $1.15 \times 10^{-31}$ |
| Interaction Between D3 and Cycloserine | 3189.2 | 6 | 531.5 | 84.2 | $6.28 \times 10^{-15}$ |
| Error | 151.4 | 24 | 6.3 | | |
| Total | 70286 | 35 | | | |

**Table S3 | List of bacterial strains used in this study**

| Strain designation | Host/parental strain | Relevant genotype/plasmid | Description | Source |
|---|---|---|---|---|
| MG1655 | | | Wild-type *E. coli* K-12 | Lab stock |
| LE392 | | $sup^E$ $sup^F$ | Nonsense suppressor | Lab stock |
| PAO1 | | | Wild-type *P. aeruginosa* | Lab stock |
| LZ613 | LE392 | (λ*cI857 bor::Kan*$^R$) | Lysogenic for lambda | Lab stock |
| LZ2001 | MG1655 | [pZS*24-$P_{FtsKi}$-*tetR-mNeonGreen*] | Phage DNA reporter strain | Lab stock |
| LZ2004 | LZ613 | [pZS*34-$P_{FtsKi}$-*tetR-mNeonGreen*] | Lambda lysogen carrying the DNA reporter plasmid | Lab stock |
| RY33001 | MG1655 | *manX::Kan*$^R$ | | Ry Young |
| RY33002 | MG1655 | *manY::Kan*$^R$ | | Ry Young |
| RY33003 | MG1655 | *manZ::Kan*$^R$ | | Ry Young |
| LZ3525 | RY33001 | [pZS*24-$P_{ftsKi}$-*tetR-mNeonGreen*] | | This work |
| LZ3526 | RY33002 | [pZS*34-$P_{ftsKi}$-*tetR-mNeonGreen*] | | This work |
| LZ3527 | RY33003 | [pZS*34-$P_{ftsKi}$-*tetR-mNeonGreen*] | | This work |
| PL15 | RP437 | *fhuA31* Δ*motAB*, *fliC*$^{[sticky]}$ | Sticky-flagella strain used for imaging live cells in suspension | Pushkar Lele |
| LZ3543 | PL15 | *mrcB:: Kan*$^R$ *fhuA*$^+$ | *fhuA* revertant version of PL15 for T5 infections | This work |
| JW0145 | BW25113 | *mrcB:: Kan*$^R$ | Keio collection *mrcB* knockout strain | Jason Gill |
| KC427 | MG1655 | *lptD4213 carB::Tn10* | Outer membrane hyperpermeable mutant (*imp4213*) | K.C. Huang |

**Table S4 | List of bacteriophages used in this study**

| Phage Designation | Relevant Genotype | Description | Source |
|---|---|---|---|
| λLZ613 | λ*cI857 bor::Kan$^R$* | "Wild-type" lambda with kanamycin resistance gene | Lab stock |
| λLZ2002 | λ*D- mTurqoise2 cI857-mKO2 bor::48×tetO- Cm$^R$* | Three-color DNA/cell-fate reporter phage | Lab stock |
| λLZ770 | λ*D- mTurqoise2 Pam80 bor::Cm$^R$* | Replication-deficient (*P* amber mutant) variant of fluorescent lambda. Propagate on the suppressor strain LE392. | Lab stock |
| T4 | | Wild-type T4 | Ry Young |
| T5 | | Wild-type T5 | Ry Young |
| T7 | | Wild-type T7 | Ry Young |
| D3 | | Pseudomonas LPS-specific Lambdoid phage | Alan Davidson |
| DMS3 | | Pseudomonas Mu-like (transposable) siphophage | Alan Davidson |

### Supplementary figures

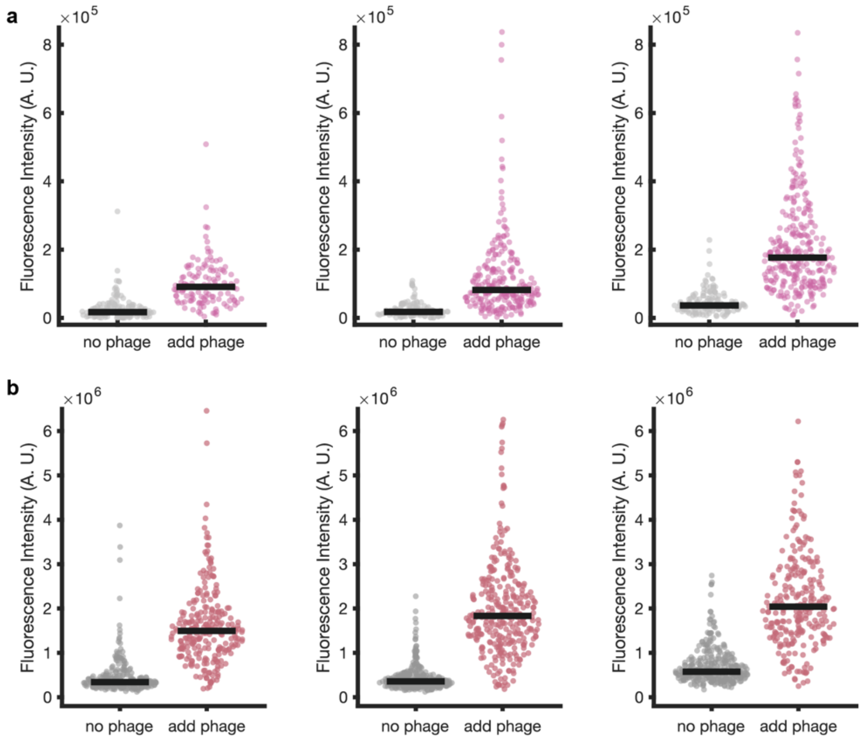


**Supplementary Fig. 1 | Individual replicates for quantifying phage λLZ613-induced small molecule accumulations.**

Three biological replicates for the comparison between phage and negative control for daunorubicin **a**), and propidium iodide **b**) at an MOI of 5.

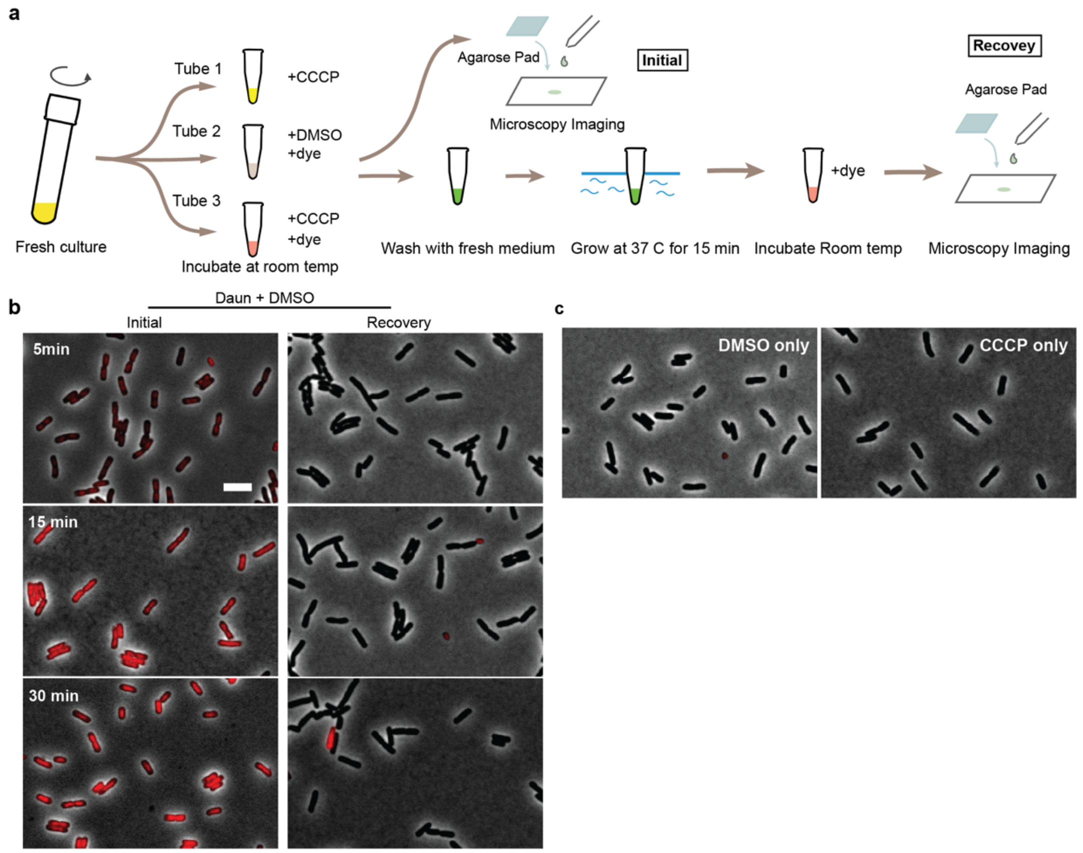


**Supplementary Fig. 2 | The effect of energy state on changing the cell permeability**

**a**) Schematic diagram of examining the transiency and reversibility of membrane potential disruption by energy poison CCCP.
**b, c**) Cells were able to recover from daunorubicin accumulation and resume normal growth, regardless of CCCP treatment duration. The control groups **c**) show no background level of daunorubicin accumulation when treated with DMSO (the solvent for CCCP), and CCCP itself does not exhibit autofluorescence within the mCherry spectrum.

Scale bars = 5 μm.

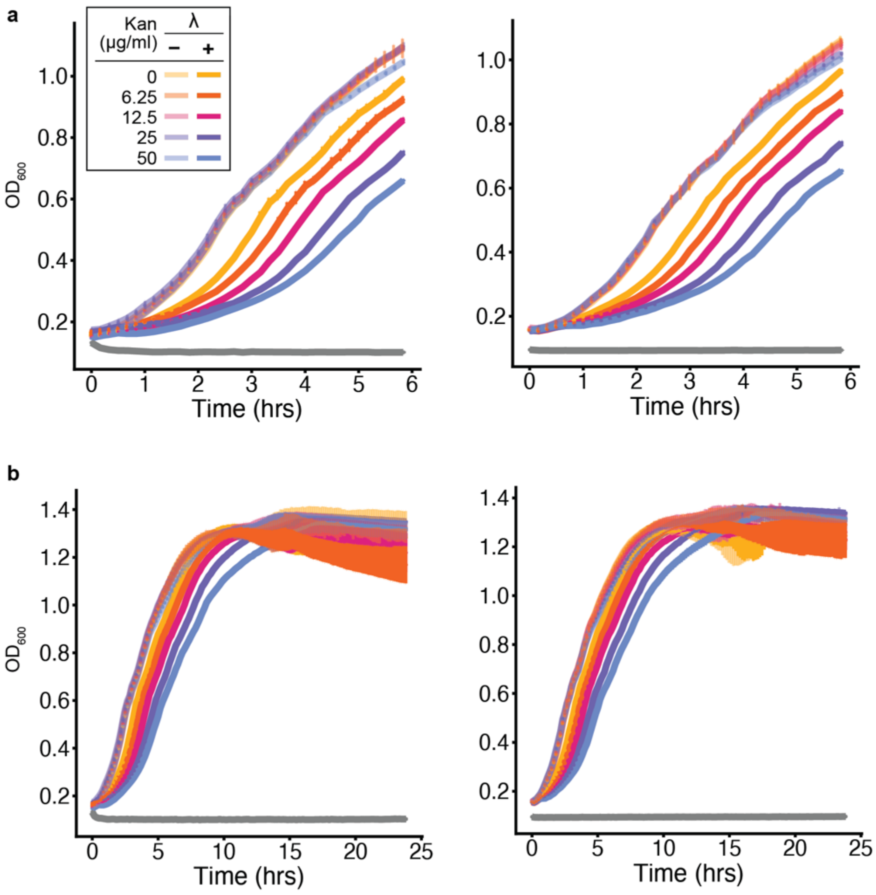


**Supplementary Fig. 3 | Additional biological replicates of the λ-kanamycin combination treatment in the lysogen.**

**a**) Growth curve measurements using the same experimental setup as in **Fig. 5b**. Wildtype phage λLZ613 added (MOI = 5) in combination with kanamycin at various concentrations resulted in an initial reduction in lysogen growth (up to ~10 hours), after which growth curves converged (**b**, 24 hours). Since no further phage replication or production occurs, it is reasonable to observe the effect only at early time points. See **Extended Data Fig. 8d** for experiments in which phages were introduced multiple times.

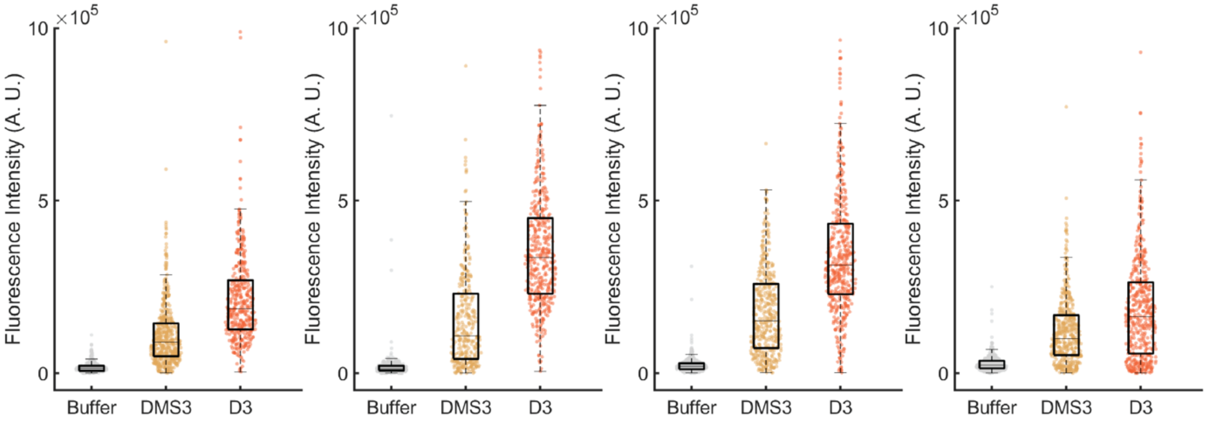


**Supplementary Fig. 4 | Individual replicates for quantifying phage DMS3- and D3-induced daunorubicin accumulations.**

Four biological replicates confirmed that DMS3- and D3-infected *P. aeruginosa* PAO1 cells show a significant increase in daunorubicin accumulation at 10 min post-infection.

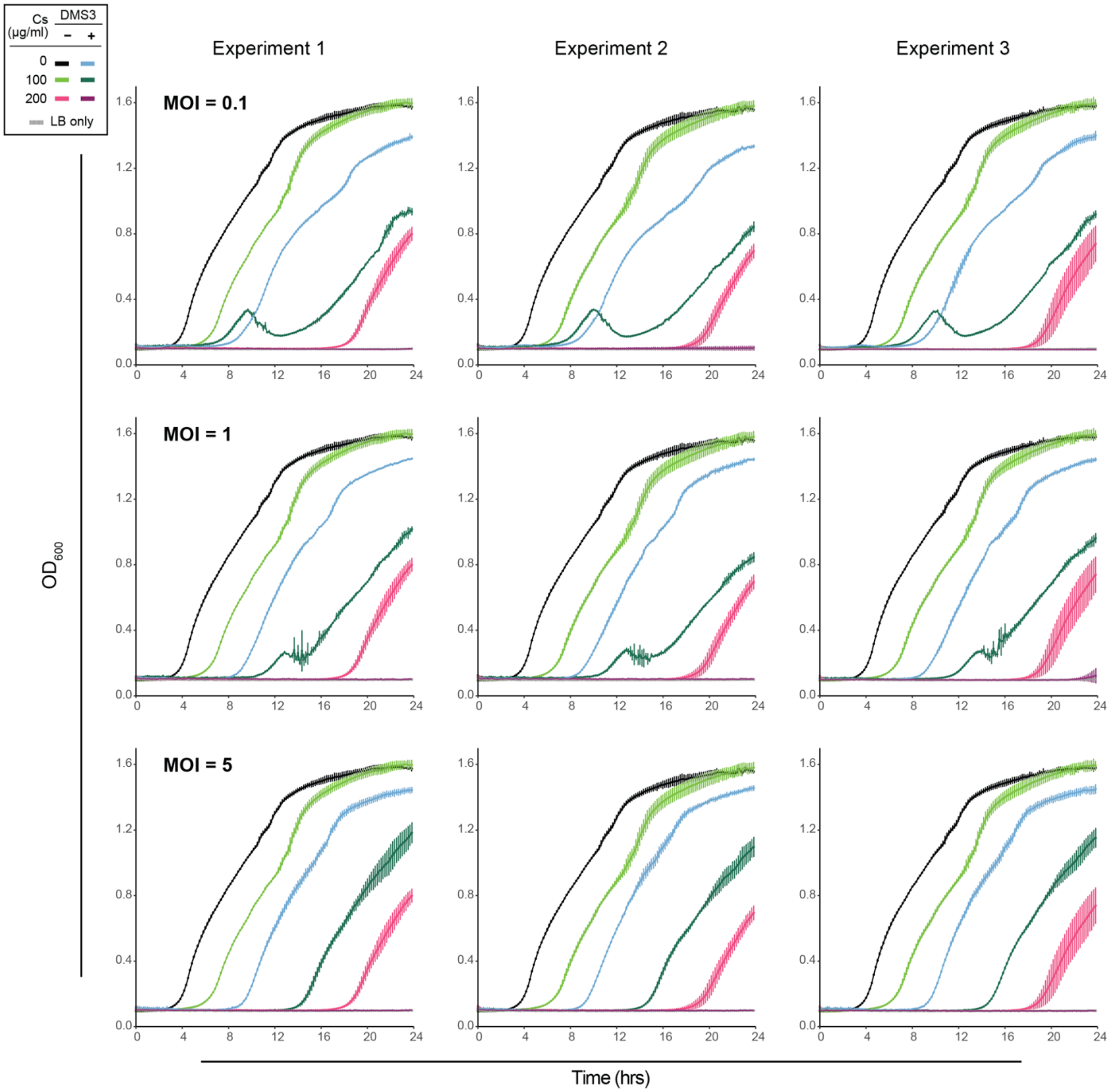


**Supplementary Fig. 5 | Individual biological replicates of DMS3-cycloserine synergy.**

Three biological replicates of growth curves of *P. aeruginosa* cultures treated with phage DMS3 and the antibiotic cycloserine at various concentrations. One of the biological replicates at high phage concentration (MOI = 5) was used in the main figure (**Fig. 5e**). Each biological replicate (one 96-well plate) contained three technical replicates (three wells) for each treatment combination. All MOI–antibiotic concentration combinations showed synergistic killing effects.

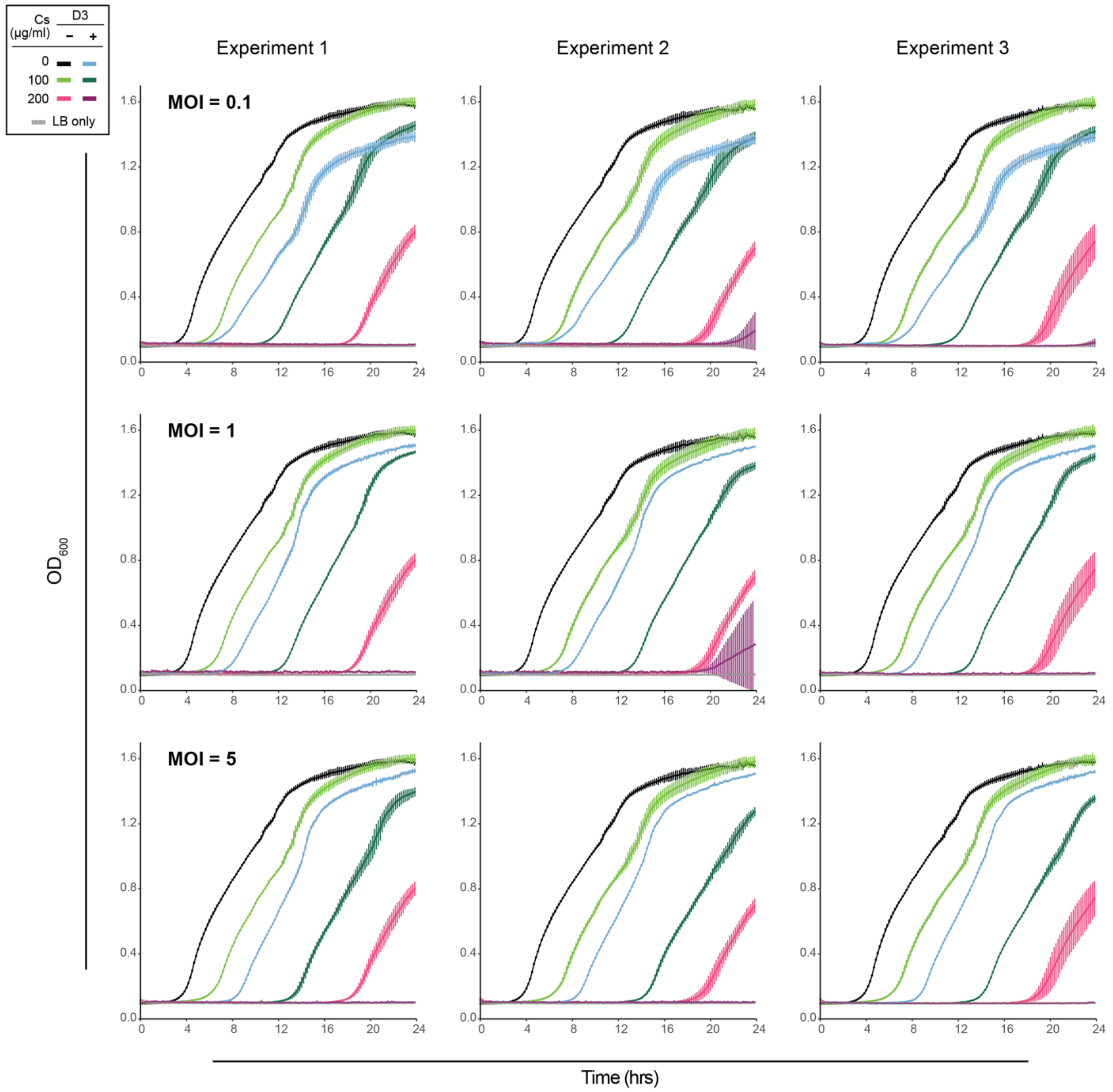


**Supplementary Fig. 6 | Individual biological replicates of D3-cycloserine synergy.**

Three biological replicates of growth curves of *P. aeruginosa* cultures treated with phage D3 and cycloserine at various concentrations. One of the biological replicates at high phage concentration (MOI = 5) was used in the main figure (**Fig. 5e**). The experimental design was the same as described above in **Supplementary Fig. 5**.
